# Structure of full-length APOBEC3B bound to EBV BORF2 reveals coordinated neutralization of a cancer-associated mutator

**DOI:** 10.64898/2026.06.18.733201

**Authors:** William Fried, Ziyuan Li, Connor Qiu, Lindsey Whalen, Hanjing Yang, Xiaojiang S. Chen

## Abstract

The double-domain APOBEC3B (A3B) contains an N-terminal non-catalytic and C-terminal active cytidine deaminase domains (NTD and CTD). A3B restricts viral infection yet is also a major endogenous mutator in human cancer, driving tumor evolution and drug resistance. A previous study showed that the Epstein-Barr virus (EBV) ribonucleotide reductase large subunit BORF2 binds the active site of the A3B CTD and neutralizes the activity of the isolated catalytic domain. However, the three-dimensional organization of full-length A3B (fl-A3B) and the contribution of A3B NTD to BORF2-mediated neutralization have remained unclear. Here, we report a 2.77 Å cryo-electron microscopy structure of fl-A3B in complex with EBV BORF2. The structure reveals an elongated hetero-octamer comprising two A3B_2_/BORF2₂ tetramers linked through alternating canonical and non-canonical BORF2 dimer interfaces. Full-length A3B adopts a defined NTD–CTD arrangement distinct from previously characterized APOBEC3G conformations. The A3B CTD binds BORF2 and occludes access to the catalytic site, while the A3B NTD simultaneously contacts a second BORF2 subunit, stabilizing a previously uncharacterized BORF2 canonical dimer interface required for higher-order assembly. Disruption of either BORF2 dimer interface prevents formation of filamentous A3B-BORF2 assemblies. These findings establish the structural architecture of full-length A3B and reveal how dual-domain engagement by A3B promotes higher-order BORF2 assembly, providing a structural framework for understanding EBV-mediated neutralization of an antiviral and cancer-associated mutator.

## Introduction

The APOBEC3 (A3) family of cytidine deaminases constitutes a central component of intrinsic immunity, restricting retroviruses, endogenous retroelements, and DNA viruses through cytosine deamination of single-stranded DNA ^1–8^. Among the seven human A3 enzymes, there are four A3 members, A3B, A3D, A3F, and A3G, containing tandem cytidine deaminase domains ^9,10^. In these double-domain A3s, catalytic activity resides in the C-terminal domain (CTD or CD2), whereas the N-terminal domain (NTD or CD1) is catalytically inactive but critically regulates nucleic acid binding, oligomerization, and enzymatic activity ^11–17^.

APOBEC3B (A3B) is unique among A3 enzymes in being constitutively nuclear ^18^, positioning it to act on both viral genomes and host DNA. While A3B contributes to antiviral defense, it has also emerged as a major endogenous mutator in human cancer ^19–22^. Elevated A3B expression correlates with cytosine mutational signatures enriched in 5′-TC motifs in multiple tumor types, with a preference for 5’-RTC contexts ^23–25^. A3B overexpression is associated with increased tumor heterogeneity, poor prognosis, and enhanced tumor evolution ^23,26–30^. Thus, A3B functions at the intersection of innate immunity and genome instability.

Despite its biological significance, the structural basis of A3B regulation remains incompletely understood. High-resolution crystal structures of the A3B NTD and CTD alone have been reported ^16,31–33^. However, full-length structures of double-domain APOBEC3s have been difficult to obtain due to poor solubility, aggregation, and RNA-associated oligomerization. Consequently, the three-dimensional organization of the NTD and CTD of intact fl-A3B remained unresolved, and it remains elusive as to how the two domains coordinate its biological activities, including oligomerization, RNA binding, enzymatic activity, and other cellular functions.

This gap is particularly important in the context of viral immune evasion. Epstein-Barr virus (EBV), a ubiquitous human gamma-herpesvirus and etiological agent of infectious mononucleosis and multiple malignancies, replicates its genome in the nucleus during lytic infection ^34–38^. EBV genomes are susceptible to A3B-mediated deamination in the absence of viral countermeasures, resulting in hypermutation, reduced viral titers, and diminished infectivity ^39^. To preserve viral genome integrity, EBV encodes a multifunctional antagonist of A3B: the large subunit of its ribonucleotide reductase (RNR), BORF2. BORF2 directly binds A3B, inhibits its DNA cytosine deaminase activity, and relocalizes A3B from the nucleus to cytoplasmic aggregates ^39–42^. Previous structural studies demonstrated how BORF2 engages the isolated catalytic CTD of A3B ^42^, but the detailed mechanisms by which EBV BORF2 recognizes fl-A3B to neutralize it remains unknown.

Determining the structure of fl-A3B is therefore essential for understanding how this enzyme functions at the intersection of antiviral immunity and cancer mutagenesis. Here, we report two cryo-EM structures of human fl-A3B in complex with EBV BORF2. These structures establish the domain architecture of a double-domain APOBEC3 enzyme and reveal how the N-terminal and C-terminal domains cooperate to form an elongated hetero-octamer that can extend into a filamentous assembly.

Our findings establish the structural architecture of full-length A3B and reveal how dual-domain engagement by A3B promotes higher-order BORF2 assembly, suggesting a potential structural link between antiviral antagonism and viral ribonucleotide reductase organization.

## Results

### Cryo-EM structure of Full-Length A3B bound to BORF2

To better understand the functional roles of A3B in protecting the cell from viral infections and cancer mutagenesis, we solved a structure of a hetero-octameric complex composed of four copies of a full-length A3B mutant (fl-A3Bm) and four copies of a truncated BORF2 construct (1-739 aa) (Fig 1a). Our fl-A3Bm construct (see Supplementary Fig. 1a, b), reduced aggregation and improved solubility when expressed as a MBP fusion, allowing for purification from *E. coli.* The BORF2 construct (1-739 aa, referred to as BORF2 hereafter) comprises the visible region of the previously solved structure of a A3B CTD/BORF2 complex ^42^ which suggests that the non-visible region is likely disordered or flexible. An N-terminal MBP tag and C-terminal His-tag was added for aiding in the purification of the BORF2 protein.

**Fig. 1.**
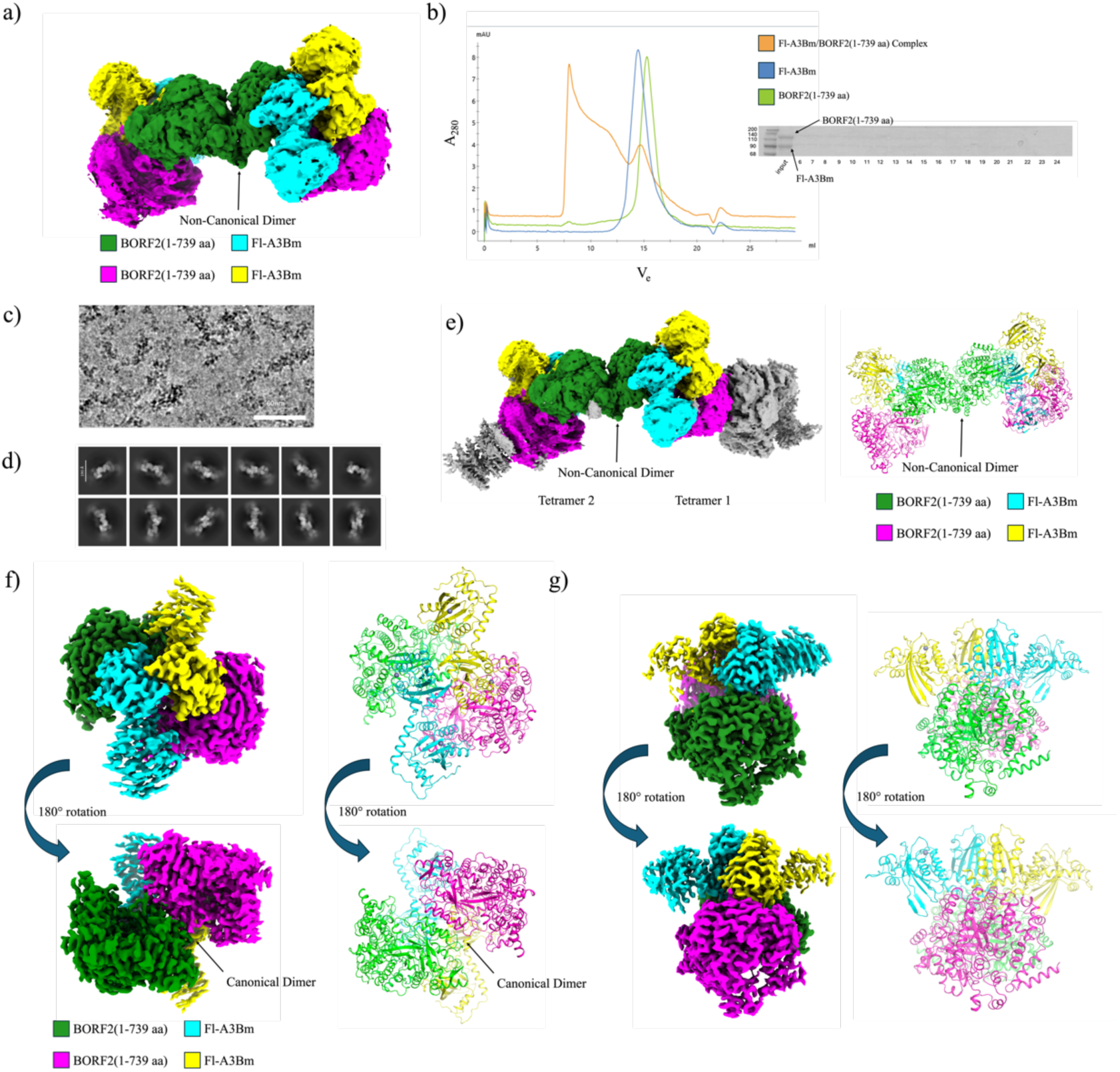
Protein purification and the overall cryo-EM structure of full-length A3B/BORF2 complex. **a)** Cryo-EM density map of the hetero-octameric fl-A3Bm/BORF2 complex used for structure determination. The complex consists of four fl-A3Bm molecules and four BORF2 (1–739 aa) subunits. **b)** Biochemical characterization of complex formation. Left, overlaid SEC profiles of fl-A3Bm/BORF2 complex (orange), fl-A3Bm alone (blue), and BORF2 (1–739 aa) alone (green). Right, SDS–PAGE analysis of the purified fl-A3Bm/BORF2 complex. **c)** Representative cryo-EM micrograph displaying multimeric particles and elongated filamentous assemblies formed by the fl-A3Bm/BORF2 complex. **d)** Selected 2D class averages showing major particle orientations used for 3D reconstruction. **e)** Left, cryo-EM density map of the fl-A3Bm/BORF2 octamer highlighting additional density corresponding to neighboring BORF2 subunits consistent with higher-order assembly. Right, ribbon representation of the octamer showing two tetramers linked through BORF2 non-canonical dimer interfaces in the middle. **f).** Sharpened cryo-EM map and corresponding ribbon structure model of one fl-A3Bm₂/BORF2₂ tetramer, shown after 180° rotation to illustrate domain organization. **g).** Sharpened cryo-EM density and ribbon structure model highlighting fl-A3Bm dimer contacts across the BORF2 canonical dimer interface, shown in two orientations (180° rotation).

When the mixture of fl-A3Bm and BORF2 proteins at a 2:1 ratio was analyzed in Superpose 6 analytical column chromatography, broad peaks of large oligomers were formed, which eluted earlier than from peaks of fl-A3Bm (MWt ∼88 KDa for a monomer) and BORF2 (∼127 kDa for a monomer). The individual peaks of fl-A3Bm eluted to a MWt between a trimer and tetramer while BORF2 eluted at a position corresponding to the MWt of a dimer (Fig. 1b). The broad peaks of the oligomers have a wide spectrum of molecular masses, ranging from roughly 400 kDa in the first peak to 5,000 kDa or more in the peak near and at the void volume, suggesting a mixture of different sizes of oligomers of the fl-A3Bm/BORF2 complex. Under cryo-EM, this protein mixture revealed protein aggregates, individual filaments of various length (Fig. 1c), as well as individual particles (Fig. 1d).

3D cryo-EM reconstruction of the single particles of the fl-A3Bm/BORF2 complex resulted a hetero-octameric complex structure to a resolution of 2.77 Å (Fig 1e; Table 1; Supplementary Fig. 2,3,4). The octameric complex can be considered as the dimer of two fl-A3Bm/BORF2 tetrameric complexes that dimerizes through the BORF2 non-canonical dimer interface (Fig. 1e), an interface that has been characterized earlier ^42^. Each tetrameric complex contains two fl-A3Bm molecules binding to two copies of BORF2 across a newly observed dimer interface of BORF2 (Fig. 1f), which is previously hypothesized and termed as the BORF2 canonical dimer interface ^42^. This newly identified BORF2-BORF2 interface is found to be stabilized by multiple interactions with both the N- and C-terminal domains (NTD and CTD) of two fl-A3Bm molecules that also show contacts with each other in an inverted orientation (Fig 1g).

**Table 1:** Cryo-EM data collection, refinement, and validation statistics.

|  | Fl-A3Bm/BORF2 hetero-octamer<br>(PDB 10VX) | Fl-A3Bm/BORF2-R39E<br>Tetramer<br>(PDB 10KX) |
| --- | --- | --- |
| **Data collection** |  |  |
| Magnification | 105,000 | 150, 000 |
| Voltage (kV) | 300 | 200 |
| Electron exposure (e<sup>-</sup>/Å<sup>2</sup>) | 65 | 50 |
| Defocus range (mm) | -0.6 to -2.0 | -0.8 to -2.0 |
| Pixel size (Å) | 0.86 | 0.73 |
| Symmetry imposed | C1 | C2 |
| Initial particle images | 8,529,358 | 1,451,270 |
| Final particle images | 563,435 | 117, 598 |
| Map resolution (Å) | 2.77 | 3.45 |
| FSC threshold | 0.143 | 0.143 |
| Map resolution range (Å) | 2.5 – 7.5 | 3.0-12.0 |
| **Refinement** |  |  |
| Model resolution (Å) | 3.2 | 3.7 |
| Map sharpening B factor (Å<sup>2</sup>) | -80.0 | -135.4 |
| No. non-hydrogen atoms | 35,813 | 16,674 |
| Protein residues | 4,402 | 2048 |
| Nucleotides | 0 | 0 |
| Ligands | 8 | 4 |
| B-factors |  |  |
| Protein | 90.84 | 94.98 |
| Nucleotide | - |  |
| Ligand | 160.35 | 161.66 |
| R.m.s. deviations |  |  |
| Bond lengths (Å) | 0.004 | 0.004 |
| Bond angles (°) | 0.993 | 0.673 |
| **Validation** |  |  |
| MolProbity score | 1.57 | 1.80 |
| Clash score | 5.03 | 7.02 |
| Poor rotamers (%) | 0.55 | 0.11 |
| **Ramachandran plot** |  |  |
| Favored (%) | 95.65 | 93.81 |
| Allowed (%) | 4.35 | 6.19 |
| Disallowed (%) | 0.0 | 0.00 |

### Structure of full-length A3B within the BORF2 complex

Previous studies have determined the atomic structures of both the individual N- or C-terminal domains of A3B ^16,31–33^, however the fl-A3B structure is lacking and it is unknown how the two cytidine deaminase domains are arranged for its functions. In this high-resolution cryo-EM structure of the fl-A3B/BORF2 complex, four fl-A3Bm subunits are present in the octamer (Fig. 2a), and structural alignment shows a high degree of similarity among the four copies (Fig. 2b), indicating the same domain arrangement between NTD and CTD of these fl-A3Bm copies. Pairwise root-mean-square deviation (RMSD) values of the superimposition of the four fl-A3Bm copies range from 0.51 to 0.63 Å, indicating near-identical conformations. Comparison with previously solved isolated A3B domains shows that the full-length structure closely resembles both the individual CTD (RMSD 0.80–1.59 Å) and NTD (RMSD 1.42 Å) structures (Fig. 2c), confirming that the mutations introduced into fl-A3Bm do not substantially alter the folding of the two domains.

**Fig. 2.**
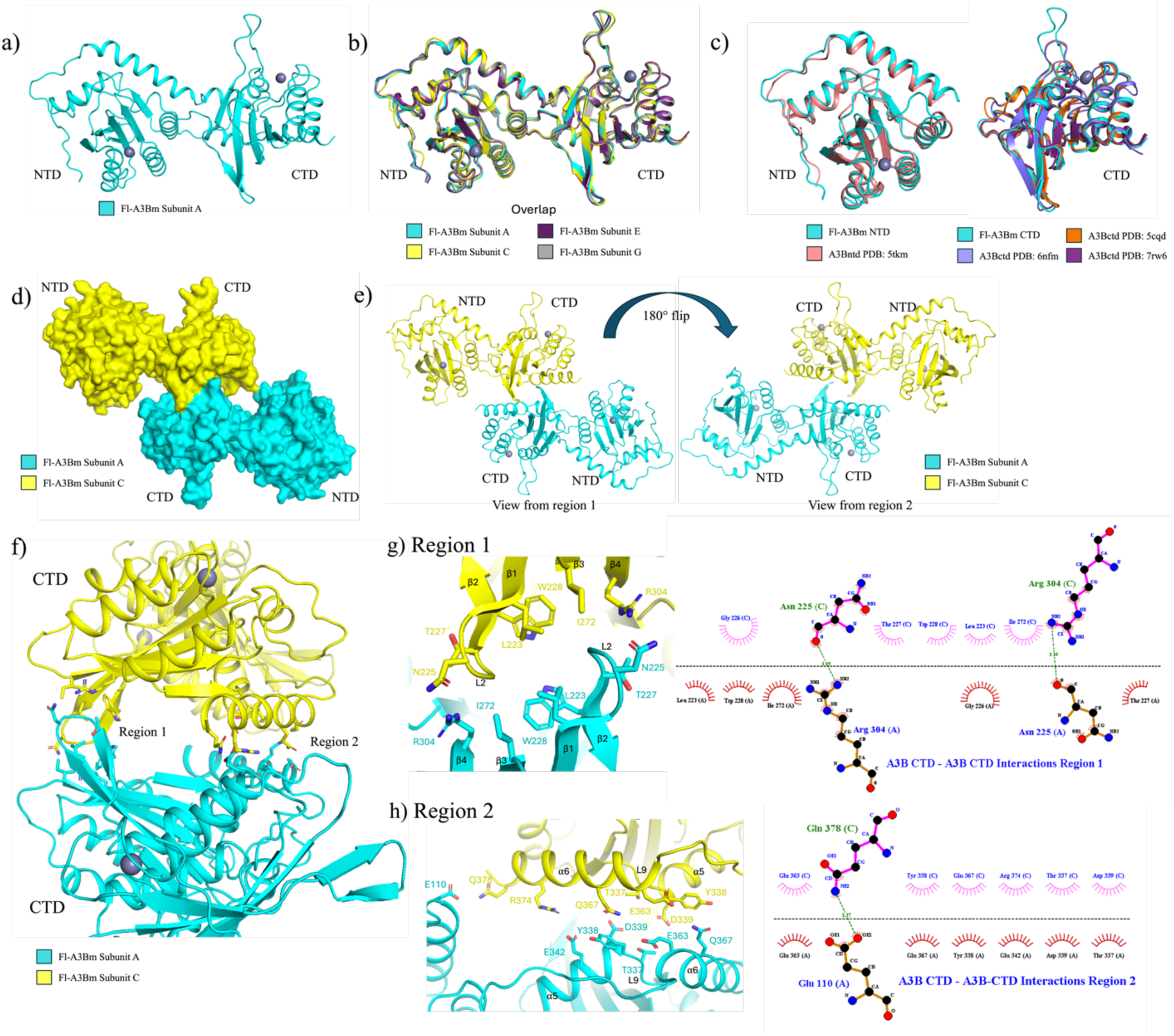
Overall structure and dimerization of full-length A3B. **a).** Ribbon representation of a single fl-A3Bm protomer extracted from the octameric complex, showing the N-terminal domain (NTD) and catalytic C-terminal domain (CTD) connected by a short linker. **b).** Structural superposition of the four fl-A3Bm protomers (subunits A, C, E, and G) within the octamer, demonstrating near-identical domain organization. **c).** Structural comparison of individual A3B domains. Left, overlay of the fl-A3Bm NTD with the previously published A3B NTD crystal structure (PDB: 5TKM). Right, overlay of the fl-A3Bm CTD with previously published A3B catalytic domain structures (PDB: 6NFM, 5CQD, and 7RW6). **d).** Surface representation of the fl-A3Bm homodimer formed within the tetrameric assembly. **e).** Ribbon representation of the fl-A3Bm homodimer shown in two orientations (including 180° rotation), highlighting the antiparallel arrangement of the catalytic domains. **f).** Close-up view of the CTD-CTD dimer interface, showing two interaction regions (Region 1 and Region 2). **g).** Detailed view of Region 1 interactions within the CTD-CTD interface, involving loop 2, β-sheet elements, and aromatic stacking contacts. **h).** Detailed view of Region 2 interactions within the CTD-CTD interface, involving loop 9 and α5-α6 helices.

The NTD and CTD of fl-A3Bm are connected by a short sequence of amino acids that links ⍺6 helix of the NTD to the ⍺1 helix of the CTD. While all four copies of A3B share similar conformations, the flexibility of the linker between NTD and CTD is apparent in the structures as the average resolution of the CTD is higher than the resolution of the NTD among the four A3B structures, possibly resulting from the much more extensive BORF2 contacts by the CTD than the NTD of A3B (Supplementary Fig. 3c).

Previously the only double-domain APOBEC member with known high-resolution structures is APOBEC3G (A3G), which reveals four different conformations with distinct domain arrangements between NTD (or CD2) and CTD (or CD2) ^43–46^. Superimposition of this fl-A3B structure with any of the A3G structures revealed this A3B structure has different NTD and CTD arrangement from the known A3G conformations (Supplementary Fig. 5a). However, aligning either the NTD or CTD of A3B with the different A3G structures shows strong structural conservation between the individual domains. Overall, in this fl-A3Bm structure the two domains are oriented in a way that show less interdomain interactions compared with A3G that displays more extensive packing interactions between the two domains in all observed A3G conformations.

Overlapping of our experimental structure with the AlphaFold Monomer v2.0-predicted A3B structure from the AlphaFold Protein Structure Database and a predicted AlphaFold3.0 structure from the Alphafold Server also revealed a major difference in the orientation of the two domains (Supplementary Fig. 5a, b) ^47–49^. Both the NTD and CTD of the AlphaFold-predicted structures and the fl-A3Bm show near identical conformations, which is expected as both the individual A3B domains have already been solved and are within the AlphaFold test database. When aligning the two structures by the NTD we see that the CTD of the AlphaFold Monomer v2.0-predicted structure is twisted about 180^°^ from where our fl-A3Bm CTD is oriented. The AlphaFold3.0 predicted structure also shows a similarly twisted conformation between the NTD and CTD domain but also shows some interdomain interactions mediated by loop 3 of the CTD.

Within the tetramer, fl-A3Bm forms a homodimer primarily mediated by CTD-CTD interactions running antiparallel orientation (Fig. 2d, e). This interface encompasses an area of about 460 Å² and consists of two distinct interaction regions (Fig. 2f). Region 1 is mediated by hydrogen bonding between A225 and R304, π-π stacking between opposing W228 residues, and additional hydrophobic contacts involving loop 2 and β-sheets 1-4 (Fig. 2g). Region 2 is mostly mediated by hydrophobic interactions and a hydrogen bond between E110 on NTD and Q378 located on CTD loop 9 and the ⍺5-⍺6 helices (Fig. 2h). These observations demonstrate that full-length A3B can dimerize through its CTD-CTD interactions.

### Characterization of the BORF2 Canonical Dimer Interface

The octameric fl-A3Bm/BORF2 structure described here is stabilized through alternating canonical and non-canonical BORF2 dimer interfaces (Fig 3a). While non-canonical dimer has been described previously ^42^, the canonical dimer had only been predicted based on homology to class I ribonucleotide reductases (RNRs), which has not been proved experimentally.

**Fig. 3.**
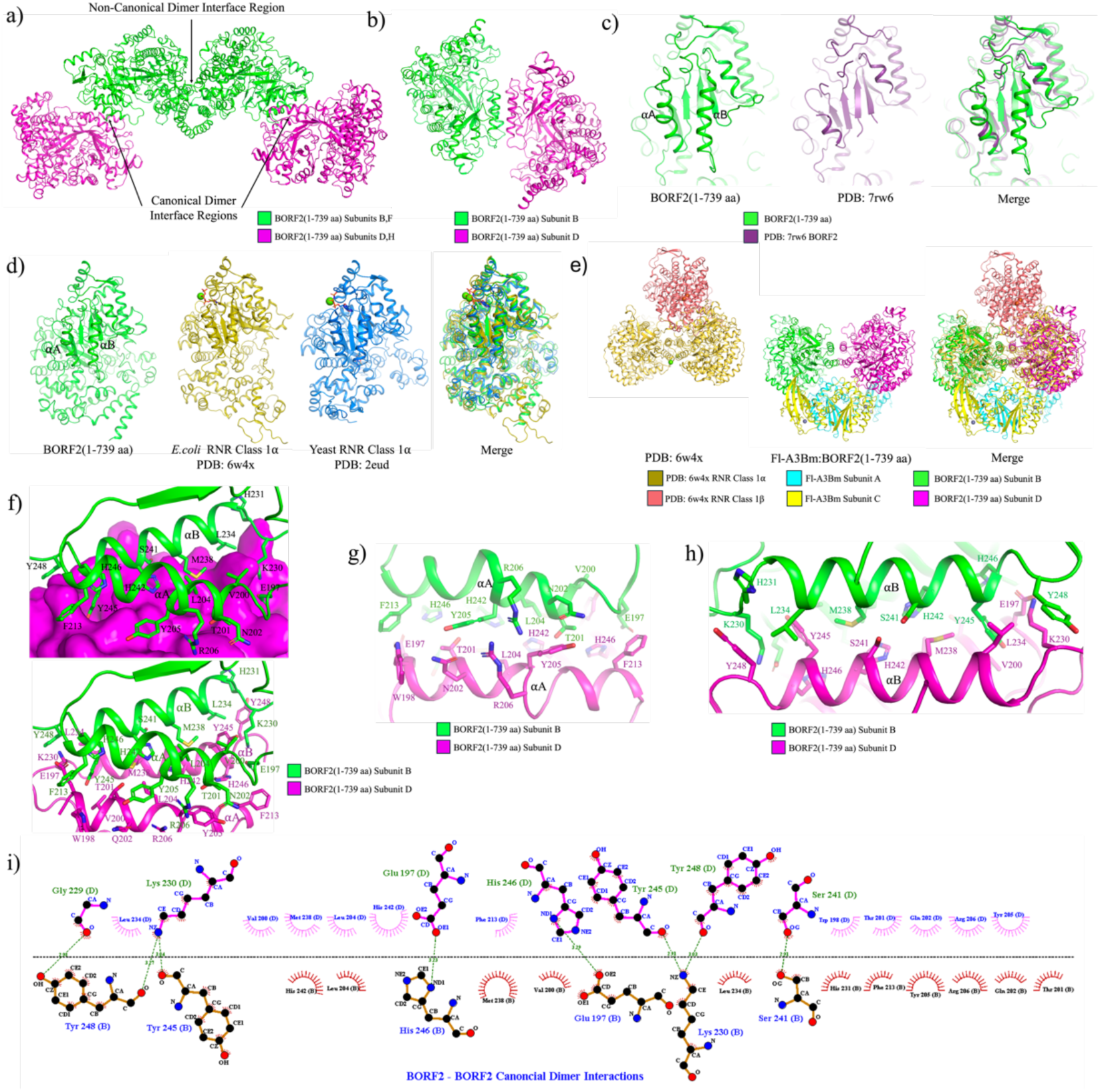
Structure of BORF2 and characterization of canonical and non-canonical dimer interfaces. **a).** Ribbon representation of the BORF2 tetramer within the fl-A3Bm/BORF2 complex, showing the alternating non-canonical and canonical dimer interfaces. The fl-A3B are removed to highlight BORF2. **b).** Detailed view of the BORF2 canonical dimer interface, showing that the αA and αB helices that constitute the four-helix bundle at this interface. **c).** Structural overlay of the BORF2 canonical dimer interface observed in this study with the previously solved BORF2 structure (PDB: 7RW6), illustrating that the αA and αB helices that constitute the four-helix bundle are unstructured in the previously reported BORF2 structure (PDB: 7RW6). **d).** Structural comparison of the BORF2 canonical dimer with cellular class I RNR α homologs from *E. coli* (PDB: 6W4X) and yeast (PDB: 2EUD), demonstrating conservation of the αA-αB helix-bundle architecture. **e).** Superposition of the fl-A3Bm/BORF2 tetramer with the *E. coli* α₂β₂ RNR complex (PDB: 6W4X), showing that A3B binding occurs on the face opposite the class I β-subunit interaction surface. **f).** Close-up view of the BORF2 canonical dimer interface showing key interacting residues and the overall four-helix αA-αB bundle organization. **g).** Detailed interactions along the αA helices of opposing BORF2 subunits, highlighting residues contributing to hydrophobic packing and hydrogen bonding. **h).** Detailed interactions along the αB helices, emphasizing symmetric insertion of Y245 and Y248 and surrounding stabilizing contacts. **i).** Ligplot of key interacting residues along BORF2 canonical dimer interface.

The canonical dimer interface is formed by two α-helices (αA and αB) from each BORF2 monomer (Fig. 3b). Notably, in contrast to previously published BORF2 structures, the region critical for interactions at this interface is disordered. In our structure, this region adopts a defined conformation, refolding into two α-helices that form the foundation of the canonical BORF2 dimer interface (Fig. 3c). These observations suggest that canonical dimerization stabilizes this region, or alternatively, that interaction-driven folding induces the structural elements required for interface formation .

Structural alignment of BORF2 with class I RNR α subunits from yeast and *E.coli* show a high degree of conservation across species, indicating that the canonical dimer interface is evolutionarily preserved (Fig. 3d). Overlapping our fl-A3Bm/BORF2 complex structure with a previously reported structure of α2β2 RNR complex from *E. coli* shows that A3B binds to the BORF2 canonical interface on the face opposite to the RNR class 1β subunit-binding surface (Fig. 3e), suggesting that A3B binding should not interfere with the 1β subunit-binding and the formation of an active α2β2 RNR complex between BORF2 and its RNR class 1β binding partner BaRF1.

The canonical dimer interface is composed of 27 residues and spans an area of 1187.4 Å^2^ , forming elaborate intermolecular bonding network (Fig 3f). It is dominated by hydrophobic interactions and hydrogen bonds distributed across the interface of a four-helix bundle that is formed between two adjacent alpha helices, ⍺A and ⍺B, of two BORF2 subunits (Fig 3f-i). A central feature of this interface is Y245 and Y248, with Y245 being symmetrically inserted into a hydrophobic pocket on the opposing BORF2 monomer, anchoring the dimer and stabilizing the interface.

### A3B stabilizes BORF2 Canonical Dimer Through its CTD- and NTD- Contacts

Within each fl-A3Bm_2_/BORF2_2_ tetramer of the octamer complex, fl-A3Bm interacts with both BORF2 monomers via its NTD and CTD across the canonical dimer interface of BORF2 (Fig. 4a). As described previously, the A3B CTD binds BORF2 primarily through interactions involving loop 1 and loop 7 around the active site of A3B CTD, encompassing a total area of about 1084.9 Å^2^ and blocking the access to the catalytic deamination site of A3B (Fig. 4b). These interactions closely resemble those observed in the earlier A3B CTD/BORF2 structure ^42^ (Fig. 4c).

**Fig. 4.**
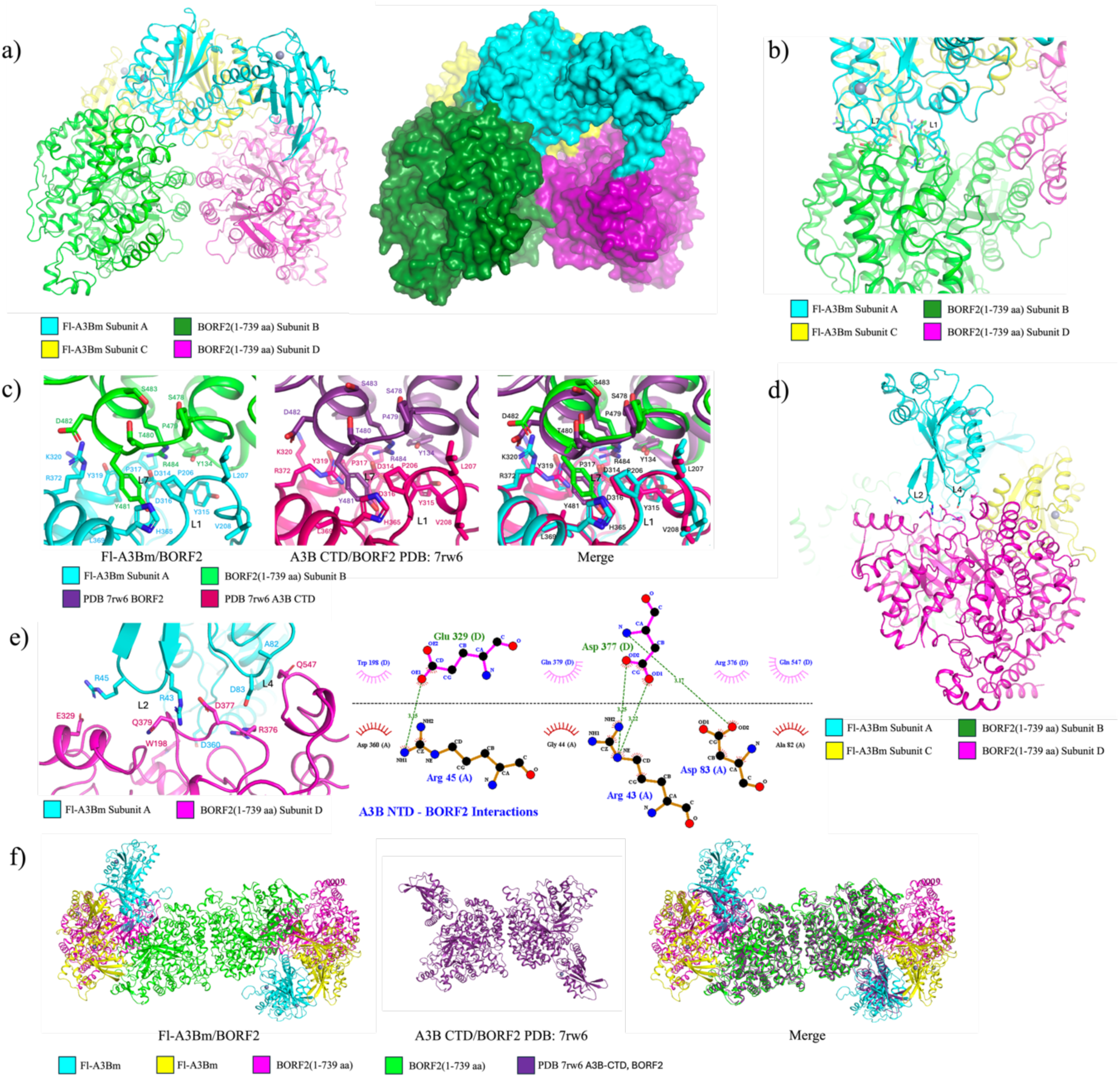
Detailed interactions between full-length A3B and BORF2 within the octamer structure. **a).** Ribbon and surface representations of a fl-A3Bm₂/BORF2₂ tetramer from the octamer showing how two A3B molecules bridge and stabilize the BORF2 canonical dimer interface. **b).** Close-up cartoon view of the fl-A3Bm CTD-BORF2 interface highlighting interactions mediated by loop 1 (L1) and loop 7 (L7) of the A3B CTD that occlude the catalytic site. **c).** Structural superposition of the CTD-BORF2 interaction observed in the full-length complex with the previously solved A3B CTD/BORF2 structure (PDB: 7RW6), demonstrating conservation of the non-canonical binding mode for the interactions with the CTD. **d).** Ribbon representation of the fl-A3Bm NTD-BORF2 interface across the canonical BORF2 dimer, showing NTD loops 2 (L2) and 4 (L4) contacting the opposing BORF2 monomer. **e).** Detailed view of the fl-A3Bm NTD-BORF2 interface highlighting hydrogen bonds and side-chain interactions that contribute to stabilization of the BORF2 canonical dimer. **f).** Structural superposition of fl-A3Bm/BORF2 hetero-octamer with previously solved A3B CTD/BORF2 structure (PDB: 7RW6), demonstrating the lack of higher-order structure in the previous structure due to missing A3B NTD domain.

In addition, we also see a previously uncharacterized interaction between the A3B NTD and the opposing BORF2 subunit of the BORF2 dimer within the tetrameric complex (Fig. 4d). This interaction is primarily mediated by loops 2 and 4 of A3B NTD and involves hydrogen bonds between A3B R45 with BORF2 E329 and A3B R43 and D83 with BORF2 D377 (Fig. 4e). Because D83 is mutated from Y83 in the fl-A3Bm construct, A3B loop 4 interactions via D83 may be partially artifactual even though a Y83 can also be modeled into this interface to form hydrogen bond and pi-charge interactions; nevertheless, these data indicate that the A3B NTD contributes to canonical dimer stabilization.

Consistent with this model of A3B NTD contributing to BORF2 canonical dimer formation and stabilization involving the NTD, the BORF2 canonical dimer was not observed in the previously reported A3B CTD/BORF2 structure that lacks A3B NTD, and which prevented the formation of higher-order fl-A3Bm/BORF2 complexes consisting of the hetero-octamer structure and its filamentous extension ^42^ (Fig. 4f). Together, these observations suggest that BORF2 canonical dimerization requires, at least in vitro, cooperative stabilization by full-length A3B, involving simultaneous interactions of A3B CTD with one BORF2 subunit and of A3B NTD with the opposing BORF2 subunit. Such interactions may be, in return, important for EBV to sequester A3B by BORF2 and shuttle A3B out of the nucleus to fend off the mutational harm of the viral genome imposed by A3B.

### Disruption of BORF2 Dimer Interfaces Prevent A3B/BORF2 Aggregation

The dual interlocking conformation of Y245 highlights the residue as critical for maintaining the canonical dimer interface of BORF2. To test the functional importance of the canonical dimer interface, we mutated Y245 to arginine, introducing a charged residue predicted to destabilize the hydrophobic core of the interface (Fig 5a). This mutant was analyzed alongside a previously characterized non-canonical dimer mutant R39E and a double mutant R39E/Y245R.

**Fig. 5.**
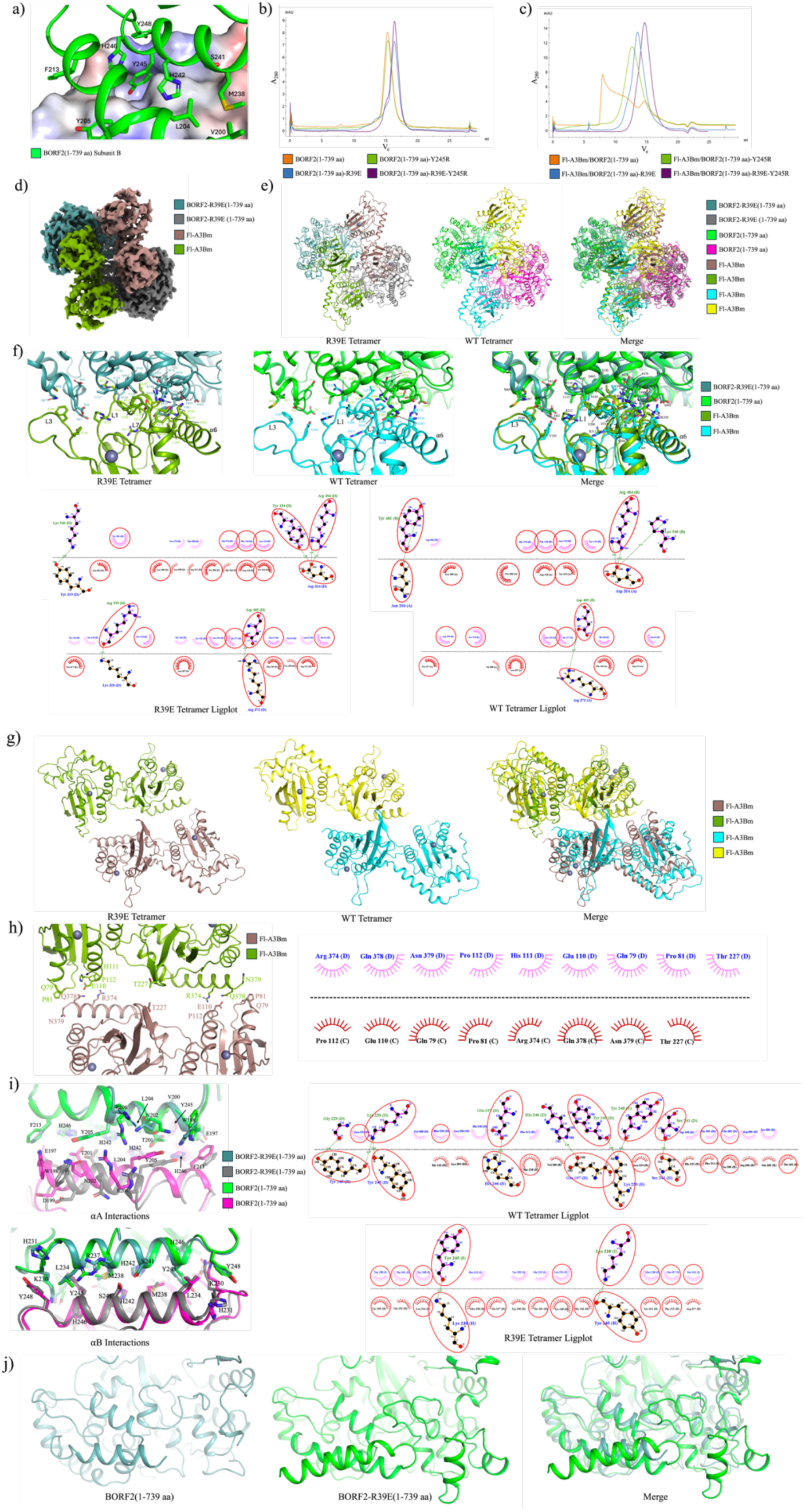
Disruption of BORF2 dimer interfaces impairs A3B-BORF2 higher-order assembly. **a).** Close-up view of the BORF2 canonical dimer interface, highlighting Y245 on one BORF2 subunit (in ribbon) inserted into a hydrophobic pocket of the opposing monomer (in surface). The Y245 interactions between the two BORF2 subunits are reciprocal. The electrostatic surface of the BORF2 subunit is colored blue for positive; red for negative; white for neutral. **b).** Overlaid size-exclusion chromatography (SEC) profiles of WT-BORF2 and BORF2 dimer-interface mutants, demonstrating altered oligomeric states upon mutation of canonical and/or non-canonical interfaces. **c).** Overlaid SEC profiles of fl-A3Bm in complex with WT BORF2 or BORF2 dimer-interface mutants, showing that disruption of either canonical or non-canonical dimerization prevents formation of high-molecular-weight aggregates. **d).** The cryo-EM map of the fl-A3Bm/BORF2-R39E tetramer particle, revealing only a tetrameric assembly in the absence of higher-order filament formation of BORF2-R39E. **e).** Structural overlay of fl-A3Bm/BORF2-R39E tetramer with the fl-A3Bm/BORF2 tetramer from the octamer complex. **f).** Top: Superposition of the A3B CTD:BORF2 interfaces from the fl-A3Bm/BORF2-R39E tetramer and the fl-A3Bm/BORF2-WT tetramer reveals tighter A3B CTD:BORF2 interactions within the fl-A3Bm/BORF2-R39E tetramer. Bottom: LigPlot comparison between the fl-A3Bm/BORF2-R39E tetramer and fl-A3Bm:BORF2-WT tetramer of key residues at the A3B CTD:BORF2 interface. **g).** Superposition of fl-A3Bm homodimers from the fl-A3Bm/BORF2-R39E tetramer and fl-A3Bm/BORF2-WT tetramer showing the shift in orientation between neighboring A3B subunits. **h)**. Detailed interactions along the interface of the fl-A3Bm homodimer form the fl-A3Bm/BORF2-R39E tetramer structure. **i).** Top left: Superposition of detailed interactions between αA helices of opposing BORF2 subunits between fl-A3Bm/BORF2-R39E tetramer and fl-A3Bm/BORF2-WT tetramer. Bottom left: Superposition of detailed interactions between αB helices of opposing BORF2 subunits between fl-A3Bm/BORF2-R39E tetramer and fl-A3Bm/BORF2 tetramer. Right: LigPlot comparison between the fl-A3Bm/BORF2-R39E tetramer and fl-A3Bm/BORF2-WT tetramer of key residues within the canonical dimer interface. **j).** Superposition of non-canonical dimer interface region between BORF2 (1-739 aa) and BORF2-R39E (1-739 aa) subunits illustrating that many of the residues comprising the non-canonical dimer interface are disordered when the non-canonical dimer is disrupted.

When analyzed by SEC in the absence of A3B, WT BORF2 elutes primarily as a dimer, whereas the BORF2 R39E mutant elutes predominantly as a monomer (Fig 5b), with a tiny shoulder peak at the dimer position. In contrast, BORF2 Y245R mutant did not alter WT-BORF2 dimerization on its own, indicating that the canonical dimer is not stably formed in the absence of A3B, at least as purified proteins *in vitro* that is free of other potential interacting components in cells.

Upon mixing with fl-A3Bm, WT BORF2 and fl-A3Bm forms large oligomers that elute across the column void volume, consistent with filament formation (Fig. 5c). Unlike the WT BORF2, all BORF2 mutants formed discrete, well-defined peaks, indicating that disruption of either the canonical or non-canonical dimer interface is sufficient to prevent non-uniform multimerization or aggregation. The BORF2 R39E/Y245R double mutant eluted the latest, consistent with a smaller complex unable to dimerize at either the canonical nor the non-canonical interfaces.

The complex of fl-A3Bm/BORF2-R39E mutant was examined by cryo-EM, which showed only single particles but no large protein aggregates and filamental structures observed in the fl-A3Bm and WT BORF2 mixture. The 3D reconstruction of the single particle images resulted in a 3.4 Å structure containing the fl-A3Bm_2_/BORF2_2_-R39E tetramer (or R39E tetramer) that corresponds to the tetramer units of the octamer (Fig. 5d, Supplementary Fig. 6,7,8, Table 1).

Aligning the R39E tetramer with the individual tetramer of the octamer complex of the WT BORF2 resulted in a RMSD of 3.425 Å highlighting that there are conformational differences between the two complexes’ structures. (Fig. 5e). The R39E tetramer has a greater interface area of 1267 Å ^2^ (average of both regions) at the A3B CTD:BORF2 interface vs the WT tetramer A3B CTD:BORF2 interface which has an area of 1079 Å^2^ (average of both regions). The larger interface area of the R39E tetramer at the A3B CTD:BORF2 interface encompasses multiple additional hydrophobic interactions and hydrogen bonds due to tighter binding of BORF2 to fl-A3Bm at this interface as well as closer engagement of the A3B loop 3 and ⍺6 helix to BORF2 (Fig. 5f).

This greater interface area appears to have pulled the neighboring fl-A3Bm CTDs away from each other, also shifting the NTD of both fl-A3Bm more towards the center of the A3B homodimer. Aligning the fl-A3Bm homodimer of both structures results in a RMSD of 6.371 Å highlighting the major shift in positioning of the fl-A3Bm homodimer in the R39E tetramer (Fig. 5g). The separation of the CTDs has weakened the interactions within the CTD-CTD dimer and has flipped the loop 2 positions of both neighboring CTDs. Within the R39E tetramer the CTD-CTD dimer is only held together by some hydrophobic interactions within the loop 2 interface whereas the region 2 interactions have been fully broken, however the new orientation of the fl-A3Bm subunits in relation to each other has resulted in additional hydrophobic interactions between the NTD and CTD domains of the opposing A3B subunits (Fig. 5h).

Within the R39E tetramer we see a reduction in size of the interface area of the canonical dimer to 1090 Å^2^ compared to an interface area of 1187.4 Å^2^ from the WT tetramer. Although aligning the two tetramer structures at the canonical dimer interface resulted in a RMSD of 0.523 Å indicating high structural homology, a shift in the positioning of ⍺1 alpha helix away from each other within the R39E tetramer resulted in a significant reduction in the number of hydrophobic interactions and hydrogen bonds across the interface (Fig. 5i). The shift within the R39E canonical dimer is likely due to the change in orientation of the fl-A3Bm homodimer which puts conformational strain on the interface.

The hydrogen bond between Y245 and K230 as well as the extensive hydrophobic interactions around Y245 are retained within the R39E structure further supporting Y245’s role in stabilizing this canonical dimer interface.

Interestingly, within the R39E tetramer we also see a lack of density around the residues that comprise the non-canonical dimer interface (Fig. 5j). This region is likely only stabilized when dimerized to the neighboring BORF2 protein or that the R39E mutation disrupts the folding of that region which then prevents the non-canonical dimer from forming. Together, these data demonstrate that both BORF2 dimer interfaces are required for higher-order oligomers of A3B/BORF2 assembly.

Notably, comparison of the fl-A3Bm/BORF2-R39E tetramer with the wild-type complex highlights the structural plasticity of multiple interfaces, including A3B:A3B, BORF2:BORF2 canonical dimers, and BORF2:A3B interactions. This interface plasticity may enable A3B/BORF2 filaments to adopt diverse conformations, facilitating their organization around the nuclear envelope to neutralize A3B, while preserving BORF2 function in nucleotide production during EBV lysogenic infection, when high levels of ribonucleotide reductase activity are required in the nucleus.

To determine whether the disruption of higher-order filamentous assembly observed *in vitro* correlates with cellular phenotypes, we evaluated the subcellular localization of wild-type BORF2-3xFlag and its dimeric interface mutants in HeLa cells co-expressing full-length A3B-Twin-Strep (**Fig. 6**). Consistent with previous reports ^39–42^, cells expressing wild-type BORF2-3xFlag and A3B-Twin-Strep forms prominent, perinuclear filamentous networks and completely sequesters host A3B out of the nuclear compartment (**Fig. 6a**).

**Fig. 6.**
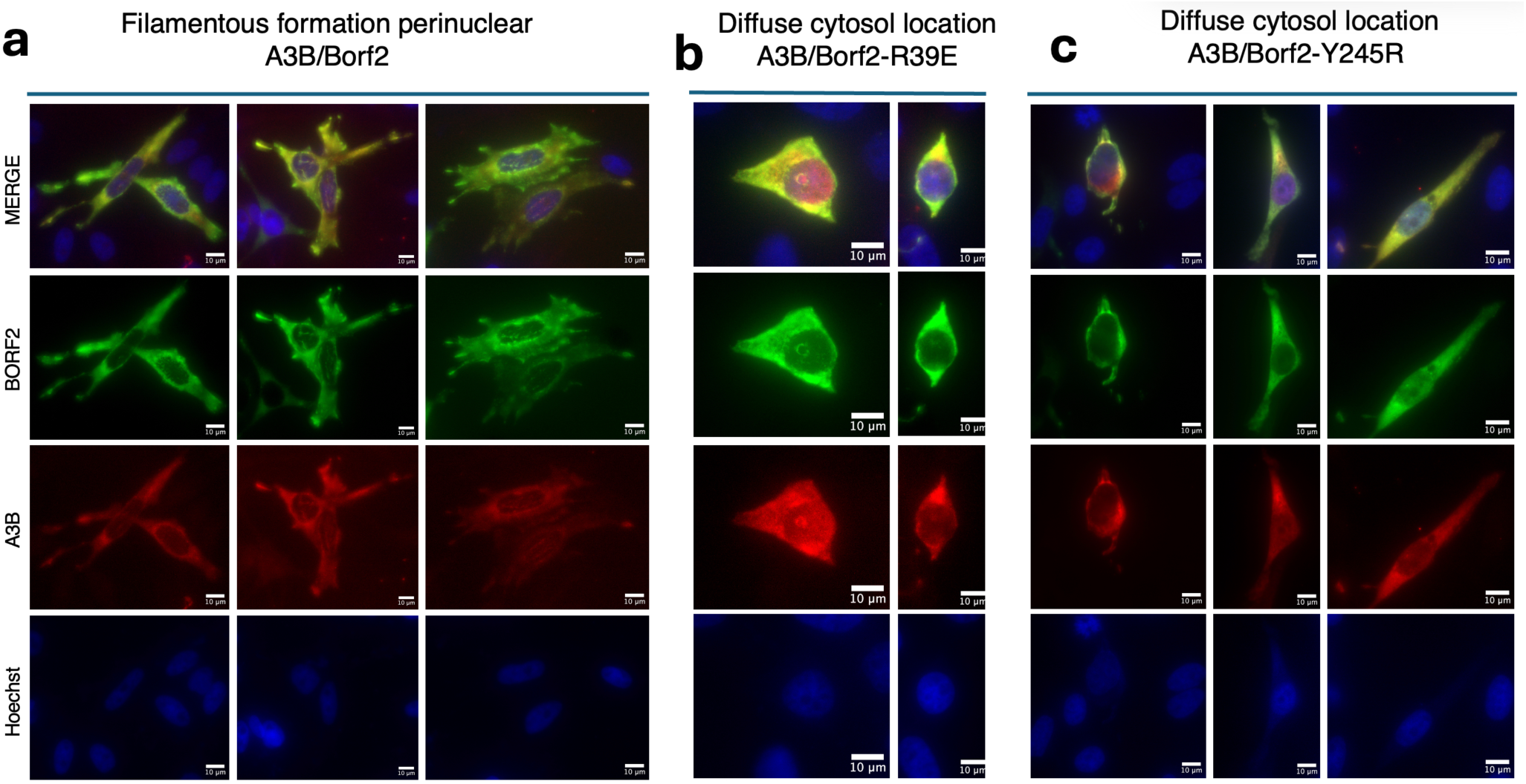
Subcellular localization of full-length A3B by BORF2 is dependent on intact dimeric interfaces of BORF2. Representative maximum intensity projection (MIP) of widefield fluorescence microscopy z-stack images of HeLa cells co-expressing full-length A3B-Twin-Strep (red) alongside BORF2-3xFlag variants (green). Nuclei are counterstained with Hoechst (blue). **(a)** Wild-type (WT) BORF2-3xFlag co-localizes with A3B-Twin-Strep, forming dense, prominent perinuclear filamentous networks. **(b)** The non-canonical dimer interface mutant BORF2-R39E-3xFlag displays a diffuse cytosolic profile and fails to assemble into networks, leaving A3B in a diffuse pan-cellular distribution. **(c)** The canonical dimer interface mutant BORF2-Y245R-3xFlag behaves similarly as BORF2-R39E-3xFlag, exhibiting a loss of higher-order networks seen for WT BORF2, yielding diffuse cytoplasmic and pan-cellular protein distributions. Scale bars = 10 µm.

In contrast, when either the non-canonical interface mutant (BORF2-R39E-3xFlag; Fig. 6b) or the canonical interface mutant (BORF2-Y245R-3xFlag; Fig. 6c) is co-expressed with A3B-Twin-Strep, this striking macromolecular filamentous networks is abolished. Instead, both mutant viral proteins display a diffuse cytosolic distribution. Under these mutant conditions, A3B exhibits a diffuse, pan-cellular profile, equilibrating across both the cytosolic and nuclear compartments. Because the R39E and Y245R mutations disrupt distinct self-association surfaces that are spatially segregated from the primary A3B-binding interface, these mutants are expected to retain baseline binding affinity for monomeric A3B. The persistent presence of A3B within the nuclear compartment alongside these mutants indicates that binary target binding alone is insufficient to clear the host mutator from the nucleus.

## Discussion

Intrinsic immunity imposes powerful selective pressure on DNA viruses that replicate in the nucleus. A3B poses a direct mutational threat to EBV genomes during lytic replication ^39^. EBV counters this threat through BORF2, which binds A3B to directly neutralize its catalytic mutational activity and relocalizes it to perinuclear bodies as an indirect neutralization of A3B by relocating A3B away from viral replicative genomic DNA in the nucleus ^39–42^. The cryo-EM structure presented here reveals that this antagonistic interaction is not a simple binary inhibition event but a higher-order architectural assembly that integrates immune neutralization of A3B with viral BORF2-mediated replication machinery organization.

### Full-length A3B structure sequestered by a viral antagonist

The hetero-octameric fl-A3Bm/BORF2 complex comprises four fl-A3Bm molecules and four BORF2 subunits. Architecturally, it can be viewed as two A3B_2_/BORF2_2_ tetramers linked through the previously characterized non-canonical BORF2 dimer interface ^42^. Within each tetramer, however, two BORF2 monomers engage through a canonical dimer interface composed of an αA:αB four-helical bundle interface. These alternating canonical and non-canonical BORF2 dimeric interfaces of the hetero-octamer complex generate a modular assembly that can extend longitudinally into higher-order filaments, as modeled in Fig. 7a. The coexistence of canonical and non-canonical interfaces establishes a structural mechanism by which A3B inhibition and RNR oligomerization occur simultaneously.

**Fig. 7.**
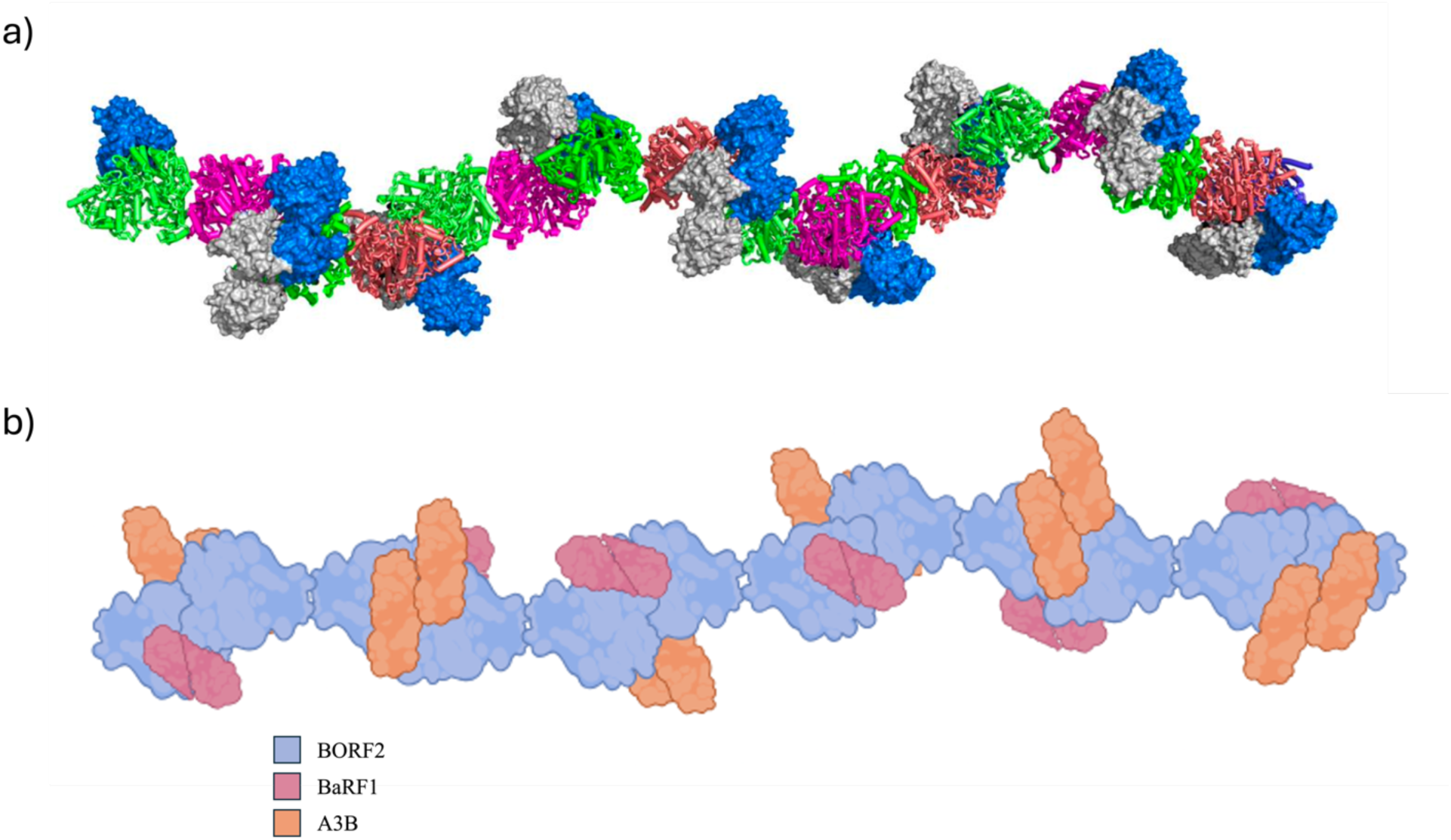
Proposed model for the EBV BORF2-mediated A3B inactivation and sequestration mechanism. **a).** Aligned model of multiple repeating fl-A3Bm/BORF2 octameric units through symmetry matrix operation based on the octamer structure, illustrating the filamentous architecture generated by alternating canonical and non-canonical BORF2 dimer interfaces. This arrangement suggests how higher-order assemblies can propagate longitudinally. **b).** Hypothetical model of the A3B/BORF2/BaRF1 complex mediated by the BORF2-A3B interactions. In this model, the A3B homodimer stabilizes the BORF2 canonical dimer interface through cooperative NTD and CTD interactions. Structural alignment with class I ribonucleotide reductases suggests that this canonical BORF2 dimer in the context of the large filamentous aggregates is compatible with association of the small subunit BaRF1 to form a putative α₂β₂ viral ribonucleotide reductase (vRNR) complex composed of BORF2 (α subunit) and BaRF1 (β subunit).

A major advance of this work is the first high-resolution structure of full-length A3B. While the isolated N-terminal (NTD) and catalytic C-terminal (CTD) domains have been previously characterized ^42^, their arrangement in the intact enzyme was unknown. In the BORF2-bound state, the NTD and CTD adopt a defined orientation connected by a short linker. Within the tetramer, two A3B molecules form an antiparallel CTD-CTD homodimer that bridges adjacent BORF2 subunits. Thus, full-length A3B becomes an architectural component of the viral assembly.

The A3B CTD binds BORF2 through loop 1 and loop 7 near the catalytic center, occluding access to substrate DNA as previously described ^42^. Unexpectedly, the A3B NTD simultaneously contacts the opposing BORF2 monomer across the canonical dimer interface. Through this dual-domain engagement, full-length A3B stabilizes the BORF2 canonical dimer, which is not stably formed in the absence of A3B. Structural alignment suggests that this canonical BORF2 dimer corresponds to the conserved RNR α-α interface and is likely compatible with interaction of the small subunit BaRF1 to form an enzymatically active α₂β₂ viral ribonucleotide reductase (RNR) complex. Although direct evidence that A3B promotes vRNR activity is currently lacking, the structural observations raise the possibility that BORF2 binding not only neutralizes A3B catalytic activity but may also stabilize a BORF2 assembly state favorable for viral DNA replication. In this view, A3B may serve not only as a restriction factor targeted for inhibition but also as a structural component that contributes to higher-order BORF2 organization.

### Canonical BORF2 dimer stabilization couples A3B neutralization to viral replication

Purified BORF2 alone predominantly forms the non-canonical dimer; stable canonical dimerization is observed only in the presence of full-length A3B. Mutation of residues within either the canonical or non-canonical interface prevents higher-order assembly and abolishes filament formation. These findings indicate that A3B binding stabilizes the canonical BORF2 dimer through cooperative interactions: the CTD engages one BORF2 monomer, the NTD contacts the second, and additional interactions across the A3B homodimer further reinforces the interface.

This cooperative stabilization suggests a potential mechanistic link between A3B neutralization and BORF2 organization. Previous studies demonstrated that BORF2 enhances EBV replication by preventing A3B-mediated hypermutation and by relocalizing A3B away from viral replication compartments ^39,40^. Notably, BORF2 does not inhibit A3B by degrading A3B but instead forms stable complexes and promotes the accumulation of A3B-containing perinuclear assemblies ^39–42^. Together with our structural observations, these findings raise the possibility that BORF2 has evolved not only to suppress A3B mutagenic activity but also to exploit A3B binding to stabilize higher-order BORF2 assemblies. Although direct effects on vRNR activity remain to be established experimentally, the coupling of A3B binding to canonical BORF2 dimer formation suggests a potential functional connection between immune neutralization and viral replication machinery organization.

### Structural basis for A3B sequestration

In cells with coexpression of BORF2 and A3B, BORF2 was previously shown to relocalizes A3B into large perinuclear bodies ^39–42^. The filament-like assemblies observed in our structural models provide the molecular basis for these aggregates, where alternating canonical and non-canonical BORF2 dimers build a longitudinal core cross-linked laterally by A3B. Our cellular imaging results validate this cooperative assembly mechanism (Fig. 6). Introducing single point mutations targeting either the four-helix bundle core (Y245R) or the non-canonical interface (R39E) completely halts filament growth, shifting BORF2 into a diffuse cytosolic phase.

Wild-type A3B is known to form high-molecular-weight complexes independently, suggesting that additional A3B-A3B interactions may further reinforce these assemblies in vivo. Thus, perinuclear body formation likely arises from a cooperative network of BORF2-BORF2, A3B-A3B, and BORF2-A3B interactions. Such multivalent assemblies may efficiently sequester A3B away from replicating viral DNA.

The filament architecture also provides a structural framework for interpreting previous cellular imaging studies. BORF2 was shown to relocalize A3B from the nucleus into large perinuclear and cytoplasmic bodies during EBV lytic replication, and similar relocalization phenotypes were subsequently observed for herpesviral BORF2 homologs ^39–41^. The elongated filamentous assemblies observed here suggest that these perinuclear bodies may not be amorphous aggregates but instead contain higher-order BORF2/A3B assemblies. Because full-length A3B contributes both CTD-mediated BORF2 binding and NTD-mediated stabilization of the canonical BORF2 dimer, the filament architecture described here may represent a structural basis for the sequestration mechanism used by EBV to neutralize A3B while simultaneously organizing BORF2 oligomers.

### Compatibility with vRNR activity

Structural alignment with bacterial and archaeal class I RNR α subunits shows strong conservation of the canonical BORF2 interface. Overlay with the *E. coli* α₂β₂ RNR complex indicates that A3B binds on the face opposite the predicted BaRF1 (β subunit) interaction surface. Therefore, A3B engagement is unlikely to sterically block formation of an active α₂β₂ vRNR complex for viral DNA replication.

Thus, our structure indicates that both the canonical and non-canonical interfaces of BORF2 can be present simultaneously within the same higher-order assembly of functional vRNR complex while sequestering A3B (Fig. 7b).

Previous studies have demonstrated that BORF2-A3B interaction enhances EBV replication by inhibiting A3B mutagenesis, relocalizing A3B away from replication compartments, and contributing to efficient lytic replication ^39–42^. Our structure suggests an additional possibility: stabilization of the canonical BORF2 dimer by full-length A3B may favor assembly of a vRNR-competent state. While this hypothesis remains to be tested directly through biochemical and virological studies, it provides a framework that reconciles the extensive and highly conserved BORF2:A3B interface, the formation of stable BORF2/A3B assemblies, and the retention rather than degradation of A3B following BORF2 binding.

### Host-pathogen conflict encoded in structure

APOBEC3 enzymes have diversified in primates under strong selective pressure from retroviruses and DNA viruses ^50–52^. The structure reported here suggests that EBV has evolved not only to neutralize A3B catalysis but also to exploit fl-A3B as a structural element that reinforces vRNR assembly. This may represent an escalation in host-pathogen conflict: a restriction factor is converted into an architectural cofactor. In this view, the A3B/BORF2 complex is both a molecular solution to immune evasion and a structural record of co-evolution.

In summary, this study establishes the structural architecture of fl-A3B and reveal how EBV integrates immune antagonism with viral replication machinery organization. By stabilizing BORF2 dimerization while blocking A3B catalytic activity, the virus converts a cancer-associated mutator and antiviral restriction factor into a structural component of its replication complex, providing a framework for understanding both viral immune evasion and the regulation of a major source of cancer genome instability.

## Methods

### Bacterial Cell Expression Constructs

An *E.coli* codon-optimized gene coding for a mutant form of human APOBEC3B (1-378)aa containing the following mutations Y13D, Y28S, F61A, Y83D, Y162D, L230S, L245K, L246H, E255A, F308Y was inserted into the pMAL expression vector along with a N-terminal MBP tag and a C-terminal 6x-His tag. The resulting MBP-fl-A3Bm-His_6_, referred to as fl-A3Bm, fusion protein was used to express protein for determining both cryo-EM structures and for SEC experiments.

A gene fragment of a truncated codon-optimized BORF2 gene (1-739)aa was inserted into a pMAL expression vector with an N-terminal MBP tag and a C-terminal 6x-His tag. The resulting MBP-BORF2(1-739)aa-His_6_, fusion protein was used to express protein for complex formation with A3B, biochemistry characterization, and the fl-A3Bm/BORF2 cryo-EM structure determination. The MBP-BORF2-R39E-His_6_, MBP-BORF2-Y245R-His_6_, and MBP-BORF2-R39E-Y245R-His_6_ mutants were generated using site-directed mutagenesis. The MBP-BORF2-R39E-His_6_ mutant was used for the fl-A3Bm/BORF2-R39E cryo-EM structure determination and for SEC experiments. The MBP-BORF2-Y245R-His_6_ and MBP-BORF2-R39E-Y245R-His_6_ mutants were used for SEC experiments.

### Mammalian Cell Expression Constructs

A mammalian codon-optimized gene coded for full-length human A3B with an AAV intron located after residue 254 was inserted into the pcDNA3.1 expression vector along with a C-terminal Twin-Strep tag. The AAV intron was included to prevent A3B-medicated self-inactivation of the vector during amplification in *E. coli*. The resulting A3B-Twin-Strep expression vector was used in immunofluorescence experiments.

A mammalian codon-optimized gene coded for full-length BORF2 was inserted into a pcDNA3.1 expression vector along with a C-terminal 3x-Flag tag. Additionally, BORF2-R39E-3x-Flag and BORF2-Y245R-3x-Flag mutants were generated using site-directed mutagenesis. Both the resulting wild-type and mutant BORF2-3x-Flag expression vectors were used in immunofluorescence experiments

### Protein Purification

For the purification of fl-A3Bm transformed *E.coli* C43(DE3) cells were cultured at 37 °C in LB medium with 1% glucose and 100 ug/mL Amp until optical density at 600 nm (OD600) reached 0.3–0.4. The growth temperature was then lowered to 16 °C and *E. coli* cells were further cultured for an additional hour. Protein expression was then induced by the addition of 0.1 mM isopropyl β-D-thiogalactopyranoside (IPTG). *E. coli* cells were cultured at 16 °C overnight and harvested by centrifugation at 3315 g for 15 minutes at 4°C. The cell pellets were then snap frozen and stored at −80 C.

The cell pellet was resuspended in lysis buffer (25 mM HEPES pH 7.4, 300 mM NaCl, 20mM MgCl2, 0.5 mM TCEP, 5% glycerol), lysed by two rounds of sonication (1 sec on, 3 sec off, 2:00 min, 75% Amp) in the presence of 1 mg of RNase A per liter of cell culture, and then incubated with 50 U/mL of Benonzase for 45 minutes at 4°C on a shaker. The lysate was then centrifuged at 25364 g for 1 hour at 4°C. The supernatant containing the His-tagged fusion protein was incubated with Ni-NTA agarose resin for 30 minutes, ran on a gravity column, and then washed by 5x 5 CV wash buffer (25 mM HEPES pH 7.4, 300 mM NaCl, 20 mM MgCl₂, 20 mM Imidazole, 0.5 mM TCEP, 5% glycerol). The trapped protein was eluted in 5x 1 CV of elution buffer (25 mM HEPES pH 7.4, 300 mM NaCl, 20 mM MgCl₂, 200 mM Imidazole, 0.5 mM TCEP, 5% glycerol).

The elutes were incubated in amylose resin for 30 minutes on a shaker then ran on a gravity column twice and washed by 3x 5 CV of lysis buffer, 5 CV of high salt buffer (25 mM HEPES pH 7.4, 1 M NaCl, 20 mM MgCl₂, 0.5 mM TCEP, 5% glycerol), and then a final 5 CV of additional lysis buffer. The sample was then eluted by 4x 5 CV of elution buffer (25 mM HEPES pH 7.4, 300 mM NaCl, 20 mM MgCl₂, 40 mM Maltose, 0.5 mM TCEP, 5% glycerol) and then each elute was mixed with equal volume of an arginine buffer (25 mM HEPES pH 7.4, 300 mM NaCl, 20 mM MgCl_2_ , 0.5 mM TCEP, 5% Glycerol, 1.0 M Arginine) to get a final arginine concentration of 0.5 M. The elutes were then concentrated to about 1 mL and then ran on an S200 prep size exclusion column equilibrated in storage buffer (25 mM HEPES pH 7.4, 300 mM NaCl, 20 mM MgCl_2_ , 0.5 mM TCEP, 5% Glycerol 0.5 M Arginine). The monomer fractions were combined and then concentrated to either 2.846 mg/mL, 3.955 mg/mL, 3.773 mg/mL, or 4.869 mg/mL. The sample was then aliquoted, snap-frozen in liquid nitrogen, and then stored at −80°C.

For the purification of BORF2(1-739)aa transformed BL21(DE3) cells were cultured at 37°C in TB medium with 100 ug/mL Amp until an optical density at 600 nm (OD600) reached 0.8-1.0 A. Protein expression was then induced by the addition of 0.5 mM IPTG. *E.coli* cells were then cultured at 18 C overnight and harvested by centrifugation at 4000g for 30 minutes at 4°C. The cell pellet was resuspended in a lysis buffer (20 mM Tris-HCl, 500 mM NaCl, 5 mM Imidazole, 5% Glycerol, 1 mM TCEP) then frozen at −80°C.

Frozen cell pellets were thawed in an ice water bath and lysed by sonication (1 sec on, 3 sec off, 2:00 min, 70% Amp) with the addition of 40 mg of lysozyme and then centrifuged at 25364 g for 1 hour at 4°C. The His-tagged fusion protein was captured by Ni-NTA agarose gravity-flow chromatography and then washed by 5x 5 CV wash buffer (20 mM Tris-HCl pH 7.4, 500 mM NaCl, 50 mM Imidazole, 5% glycerol, 1 mM TCEP). The trapped protein was then eluted via 3x 1 CV of 100 mM Imidazole elution buffer (20 mM Tris-HCl pH 7.4, 500 mM NaCl, 100 mM imidazole, 5% glycerol, 1 mM TCEP) followed by 3x 1 CV of 200 mM Imidazole elution buffer (20 mM Tris-HCl pH 7.4, 500 mM NaCl, 200 mM imidazole, 5% glycerol, 1 mM TCEP) and then concentrated to around 2 mL. The concentrated protein was then run on an S200 prep size exclusion column equilibrated with storage buffer (20 mM Tris-HCl pH 7.4, 200 mM NaCl, 5% glycerol, 1 mM TCEP). Fractions containing the eluted protein was then concentrated to 7.842 mg/mL and then aliquoted, snap frozen in liquid nitrogen and stored at −80°C.

The purification of the BORF2(1-739)aa-R39E and the BORF2(1-739)aa-Y245R constructs were initially purified as done for wild type BORF2(1-739)aa, but after the protein was eluted from the nickel resin the elutes were then incubated in amylose resin for 30 minutes on a shaker, ran on a gravity column, and washed by 5x 5 CV of storage buffer (20 mM Tris-HCl, 200 mM NaCl, 5% Glycerol, 1 mM TCEP). The trapped protein was eluted by 5x 1 CV of elution buffer (20 mM Tris-HCl, 500 mM NaCl, 200 mM Imidazole, 5% glycerol, 1 mM TCEP) and then concentrated to either 1.3 mL or 1.4 mL. The concentrated protein was then run on an S200 prep size exclusion column equilibrated with storage buffer. The fractions containing the eluted protein were then concentrated to 8.765 mg/mL for BORF2-R39E and 3.880 mg/mL for BORF2-Y245R. The concentrated protein was then aliquoted, snap frozen in liquid nitrogen and stored at −80°C.

The purification of the BORF2(1-739)aa-R39E-Y245R construct was done similarly as the purification of the other BORF2 mutant constructs with the only difference being that the protein was incubated with Ni-NTA agarose resin for 30 minutes before running on a gravity column. The protein was concentrated to 4.157 mg/mL after being run on an S200 prep and then aliquoted, snap frozen in liquid nitrogen and stored at −80°C.

### Cryo-EM Data Acquisition

Purified fl-A3Bm and truncated BORF2(1-739)aa were thawed in an ice water bath, combined at a 2:1 molar ratio of fl-A3Bm to BORF2(1-739)aa, buffer exchanged in an imaging buffer (20 mM Tris-HCl pH 7.4, 200 mM NaCl, 0.5 mM TCEP), and concentrated to a final concentration of 1.903 mg/mL. The complex was then stored in −80°C. 3uL aliquots of the fl-A3Bm/BORF2 mixture were applied to UltrAu foil R1.2/1.3 gold 300-mesh grids (Electron Microscopy Sciences). The grids were then blotted and vitrified in liquid ethane using a Vitrobot Mark IV (Thermo Fisher Scientific).

Cryo-EM data for the fl-A3Bm/BORF2 hetero-octamer complex were collected in a Titan Krios G3i (Thermo Fisher Scientific) equipped with a K3 direct electron detector and post-BioQuantum GIF energy filter (Gatan) operated at 300 kV in electron counting mode. Movies were collected at a nominal magnification of 105,000× in super-resolution mode after binning by a factor of 2, resulting in an effective pixel size of 0.86 Å. A total dose of 65 e-/Å2 per movie was used with a dose rate of approximately 15 e-/pix/sec. A total of 24,735 movies were recorded by automated data acquisition with EPU.

Cryo-EM data for the fl-A3Bm/BORF2-R39E tetramer complex were collected in Glacios (Thermo Fisher Scientific) equipped with Falcon-4 direct electron detector operated at 200 kV in electron counting mode. Movies were collected at a nominal magnification of 150,000× and a pixel size of 0.73 Å in EER format. A total dose of 50 e-/Å2 per movie was used with a dose rate of 5-6 e-/Å2/sec. 8,249 movies were recorded by automated data acquisition with EPU.

### Cryo-EM Data Processing

For the fl-A3Bm/BORF2 hetero-octamer complex a total of 24,735 movies were imported into cryoSPARC software package and were subjected to patch motion correction and CTF estimation in cryoSPARC ^53^. A total of 8,529,358 particles were picked initially, extracted, and down-sampled by a factor of 4, on which 2D classification was performed. 2,239,356 particles from 2D class averages were selected and re-extracted with full resolution. A subset of 2D class averages containing 878,874 particles was selected. 3D ab initio reconstruction was then performed to generate three initial volumes. A single class containing 578,534 particles was selected, and non-uniform refinement was performed with C1 symmetry to yield the final 2.77 Å resolution map.

For the flA3B-m/BORF2-R39E tetramer complex, a total of 8,249 movies were imported into cryoSPARC software package and were subjected to patch motion correction and CTF estimation in cryoSPARC ^53^. A total of 1,451,270 particles were picked initially, extracted, and down-sampled by a factor of 2, on which 2D classification was performed. 436,349 particles from 2D class averages were selected and re-extracted with full resolution. A subset of 2D class averages containing 242,754 particles was selected. 3D ab initio reconstruction was then performed to generate two initial volumes. A single class containing 117,598 particles was selected, and non-uniform refinement was performed with C2 symmetry to yield the final 3.45 Å resolution map.

### Model building and refinement

For the fl-A3Bm/BORF2 hetero-octamer complex an atomic model derived from the BORF2 dimer from PDB entry 7RW6 and the A3G FKL structure from PDB entry 6P40, which served as an initial placeholder for fl-A3B, was initially docked into the cryo-EM map using ChimeraX ^42,43,54^. The A3B CTD from PDB entry 5CQH and the A3B NTD from entry 5TKM was then used to replace the A3G model while preserving the initial positioning ^16,31^. After approximate placement of all components within the density map, site-specific mutations were introduced, and any required linkages between residues were modeled to generate the initial atomic model. The model was then refined using the phenix.real_space_refine module in Phenix version 1.21.2-5419 ^55^. In between refinement runs the model was further corrected manually in Coot ^56^. The final atomic model statistics were generated in Phenix and shown in Table1 ^55^. All structural figures generated with either ChimeraX or PyMol ^54^. Figures plotting interactions between protein residues were made using LigPlot^+^ ^57^.

For the fl-A3Bm/BORF2-R39E tetramer complex, an atomic model of A3B and BORF2 derived from A3B/BORF2 hetero-octamer were initially docked separately into the cryo-EM map using ChimeraX ^42,43,54^. After approximate placement of all components within the density map, site-specific mutations were introduced, and any required linkages between residues were modeled to generate the initial atomic model. The model was then refined using the phenix.real_space_refine module in Phenix version 1.21.2-5419 ^55^. In between refinement runs the model was further corrected manually in Coot ^56^. The final atomic model statistics were generated in Phenix and shown in Table1 ^55^. All structural figures generated with either ChimeraX or PyMol ^54^

### SEC analysis

For the SEC profiles fl-A3Bm, BORF2(1-739aa), and the BORF2(1-739aa) mutants were expressed and purified as described in the previous sections. For the individual protein runs, about 155 ug of fl-A3Bm or 114 ug of BORF2 was diluted in a running buffer of 20 mM Tris-HCl pH 7.4, 200 mM NaCl, 5% glycerol, 1 mM TCEP with a total sample volume of 500 uL and run on a Superose6 10/300 column equilibrated in the same running buffer.

For the SEC profiles containing the fl-A3Bm/BORF2(1-739)aa complex runs, both 155 ug of fl-A3Bm and 114 ug of BORF2(1-739)aa or BORF2(1-739)aa mutants (2:1 molar ratio) were mixed and diluted in a running buffer to equal 500 uL then incubated on ice for 30 minutes to allow the complex to form. The complex was then run on a Supercose6 10/300 column equilibrated in the same running buffer.

### Immunofluorescence Experiments

HeLa cells were maintained in Dulbecco’s modified Eagle’s medium (DMEM) supplemented with 10% fetal bovine serum (FBS). 100,000 HeLa cells were seeded into 24-well plates containing a Poly-D-Lysine (PDL) coated cover slip. After 24 hours cells were transfected with 250 ng pcDNA3.1-A3B-twin-strep and 250 ng pcDNA3.1-BORF2-3x-Flag or the listed mutant derivative constructs using Lipofectamine^TM^ 3000 following the recommended reaction conditions for a 24-well plate setup with 1.5 uL Lipofectamine^TM^ 3000 reagent. (Themo Fisher Scientific) Two days later cells were washed once with 1x phosphate-buffered saline (PBS), fixed using 4% paraformaldehyde for 15 minutes, and washed three times for 5 minutes in 1x PBS with gentle rocking. Cells were permeabilized with 0.2% Triton X-100 for 10 minutes, washed three times for 5 minutes in 1x PBS, then blocked for an hour at room temperature in 4% BSA. Cells were then incubated with primary rabbit anti-flag (Thermo Fisher Scientific, 1:500) and mouse anti-strep II (Thermo Fisher Scientific, 1:500) antibodies in 4% BSA for 1 hour at room-temperature. After washing three times for 5 minutes in 1x PBS cells were incubated with secondary goat anti-rabbit Alexa Fluor 488 (Thermo Fisher Scientific, 1:1000) and goat anti-mouse Alexa Fluor 647 (Thermo Fisher Scientific, 1:1000) antibodies in 4% BSA for 1 hour at room temperature. After washing three times for 5 minutes in 1x PBS, cell nuclei were stained with Hoechst 33342 for 10 minutes at room temperature. Cells were then washed three times for 5 minutes in 1x PBS, then the cover slips were mounted on slides using VECTASHIELD Mounting Solution (Vector Labs).

Images were collecting using a MICA Microhub (Leica Microsystems) using a 63x water-immersion objective. To capture the full three-dimensional expression profile across varying cellular volumes, automated serial optical sections (Z-stacks) were acquired over a ∼10 µm range with a fixed step size of 0.5 µm. Raw optical planes were processed using a Maximum Intensity Projection (MIP) algorithm to compress peak pixel intensities from all 21 captured layers into individual 2D composite representations using Fiji, then analyzed phenotypically.

## Data availability

The atomic models have been deposited in the PDB with accession codes: 10VX (hetero-octamer) and 10KX (hetero-tetramer). The cryo-EM maps have been deposited in the EMDB with accession codes: EMD-75497 (hetero-octamer) and EMD-75260 (hetero-tetramer).

## Acknowledgments

Electron microscopy data were collected at the Core Center of Excellence in Nano Imaging (CNI) at USC. Cryo-EM data was computed at Center for Advanced Research Computing (CARC) at USC. We thank Htet Khant for assisting the operation and maintenance of the cryo-EM facility at CNI, Tomek Osinski for assisting computing work at CARC. Fluorescence microscopy images were acquired at the Translational Imaging Center (TIC) at USC. We thank Arkadi Schwartz for technical assistance with fluorescence imaging. This work was supported by NIH grant R01AI150524 to X.S.C.

## Author Contributions

X.S.C. conceived and supervised the study. W.F. and Z.L. performed cryo-EM sample preparation, data collection and analysis, structure determination, and model refinement. C.Q., L.W., and W.F. designed and generated bacterial expression constructs of BORF2 mutants. H.Y. assisted in the design of the soluble A3B mutant construct. W.F. and L.W. expressed proteins. W.F. purified proteins and performed biochemistry characterization. W.F., C.Q., L.W., H.Y. designed, performed, and analyzed the cellular fluorescent imaging data. W.F. drafted the manuscript. All authors contributed to data analysis, discussed the results, and revised the manuscript.

## Competing financial interests

The authors declare no competing interests.

## Notes

### Competing Interest Statement

The authors have declared no competing interest.

