## Supplemental figures 1-8 for "Structure of full-length APOBEC3B bound to EBV BORF2 reveals coordinated neutralization of a cancer-associated mutator"

**for**

a)

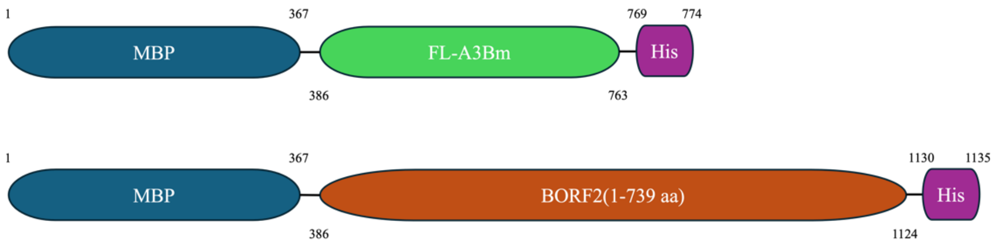


b)

```
                1                                                            60
WT-A3B          MNPQIRNPMERMYRDTFYDNFENEPILYGRSYTWLCYEVKIKRGRSNLLWDTGVFRGQVY
FL-A3Bm         MNPQIRNPMERMDRDTFYDNFENEPILSGRSYTWLCYEVKIKRGRSNLLWDTGVFRGQVY

                61                                                          120
WT-A3B          FKPQYHAEMCFLSWFCGNQLPAYKCFQITWFVSWTPCPDCVAKLAEFLSEHPNVTLTISA
FL-A3Bm         AKPQYHAEMCFLSWFCGNQLPADKCFQITWFVSWTPCPDCVAKLAEFLSEHPNVTLTISA

                121                                                         180
WT-A3B          ARLYYYWERDYRRALCRLSQAGARVTIMDYEEFAYCWENFVYNEGQQFMPWYKFDENYAF
FL-A3Bm         ARLYYYWERDYRRALCRLSQAGARVTIMDYEEFAYCWENFVDNEGQQFMPWYKFDENYAF

                181                                                         240
WT-A3B          LHRTLKEILRYLMDPDTFTFNFNNDPLVLRRRQTYLCYEVERLDNGTWVLMDQHMGFLCN
FL-A3Bm         LHRTLKEILRYLMDPDTFTFNFNNDPLVLRRRQTYLCYEVERLDNGTWVSMDQHMGFLCN

                241                                                         300
WT-A3B          EAKNLLCGFYGRHAELRFLDLVPSLQLDPAQIYRVTWFISWSPCFSWGCAGEVRAFLQEN
FL-A3Bm         EAKNKHCGFYGRHAALRFLDLVPSLQLDPAQIYRVTWFISWSPCFSWGCAGEVRAFLQEN

                301                                                         360
WT-A3B          THVRLRIFAARIYDYDPLYKEALQMLRDAGAQVSIMTYDEFEYCWDTFVYRQGCPFQPWD
FL-A3Bm         THVRLRIYAARIYDYDPLYKEALQMLRDAGAQVSIMTYDEFEYCWDTFVYRQGCPFQPWD

                361
WT-A3B          GLEEHSQALSGRLRAILQNQGN
FL-A3Bm         GLEEHSQALSGRLRAILQ----
```

**Supplementary Figure 1. Design of soluble full-length A3B mutant (fl-A3Bm) and truncated BORF2 (1-739 aa) constructs. (a)** Design of fl-A3Bm and BORF2 fusion proteins used in study. **(b)** Sequence alignment of fl-A3Bm with WT-A3B highlighting locations of mutations within fl-A3Bm.

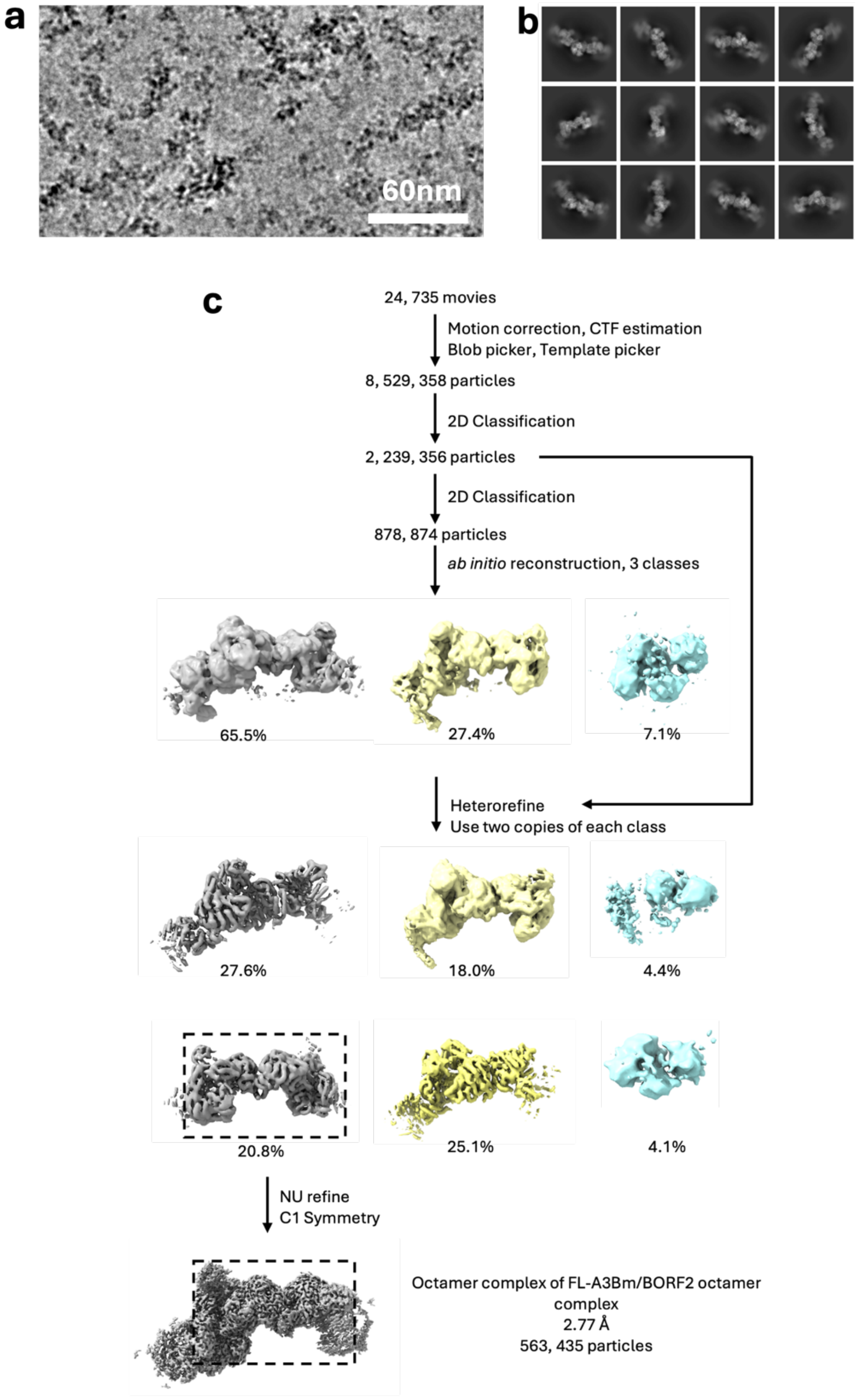


**Supplementary Figure 2. Workflow of cryo-EM 3D reconstruction of the octameric fl-A3Bm/BORF2 complex.** (**a**) Representative cryo-EM raw image, showing the filamentous feature of the complex. (**b**) Representative 2D class averages. **(c)** Cryo-EM image processing workflow. Notice the extra electron density on both ends of the octamer (boxed in dashed line), which comes from the neighboring subunits of the elongated filamentous images. However, this extra electron density, though easily detected, is not well-featured for model building.

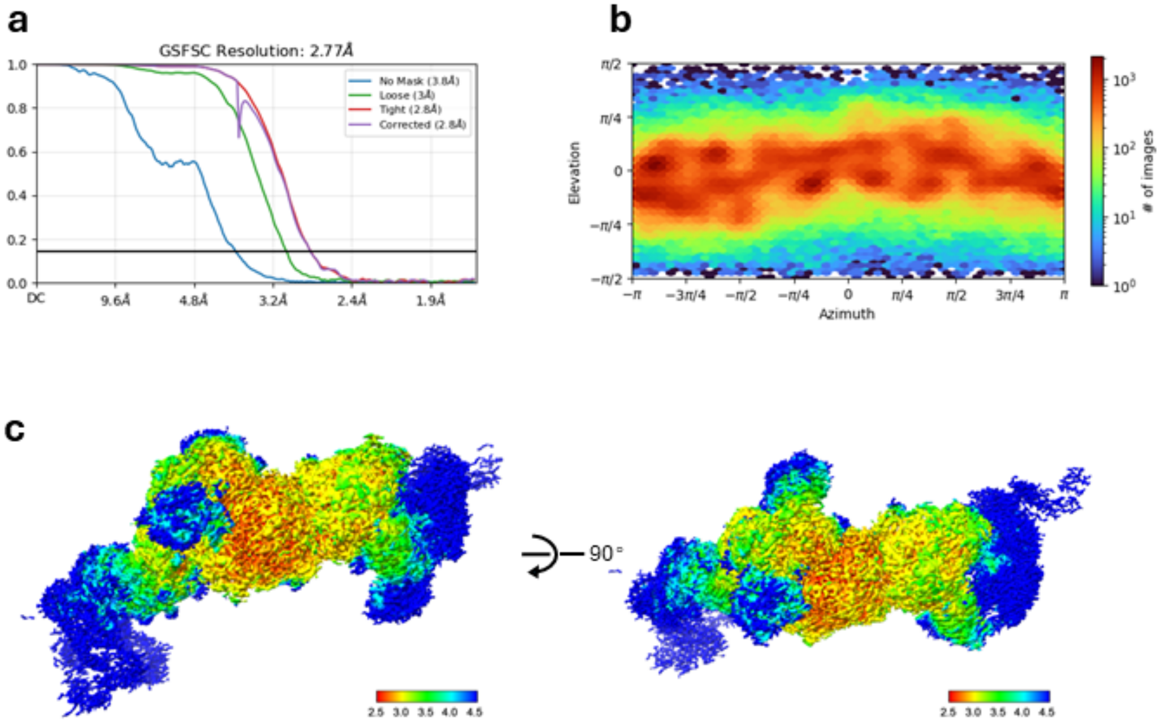


**Supplementary Figure 3. Global resolution, particle angular distribution, and local resolution estimation of the octameric fl-A3Bm/BORF2 complex.** (**a)** Global resolution estimation. (**b)**. Angular distribution plot of the particles. **(c)** Local resolution evaluation of the fl-A3Bm-BORF2 complex.

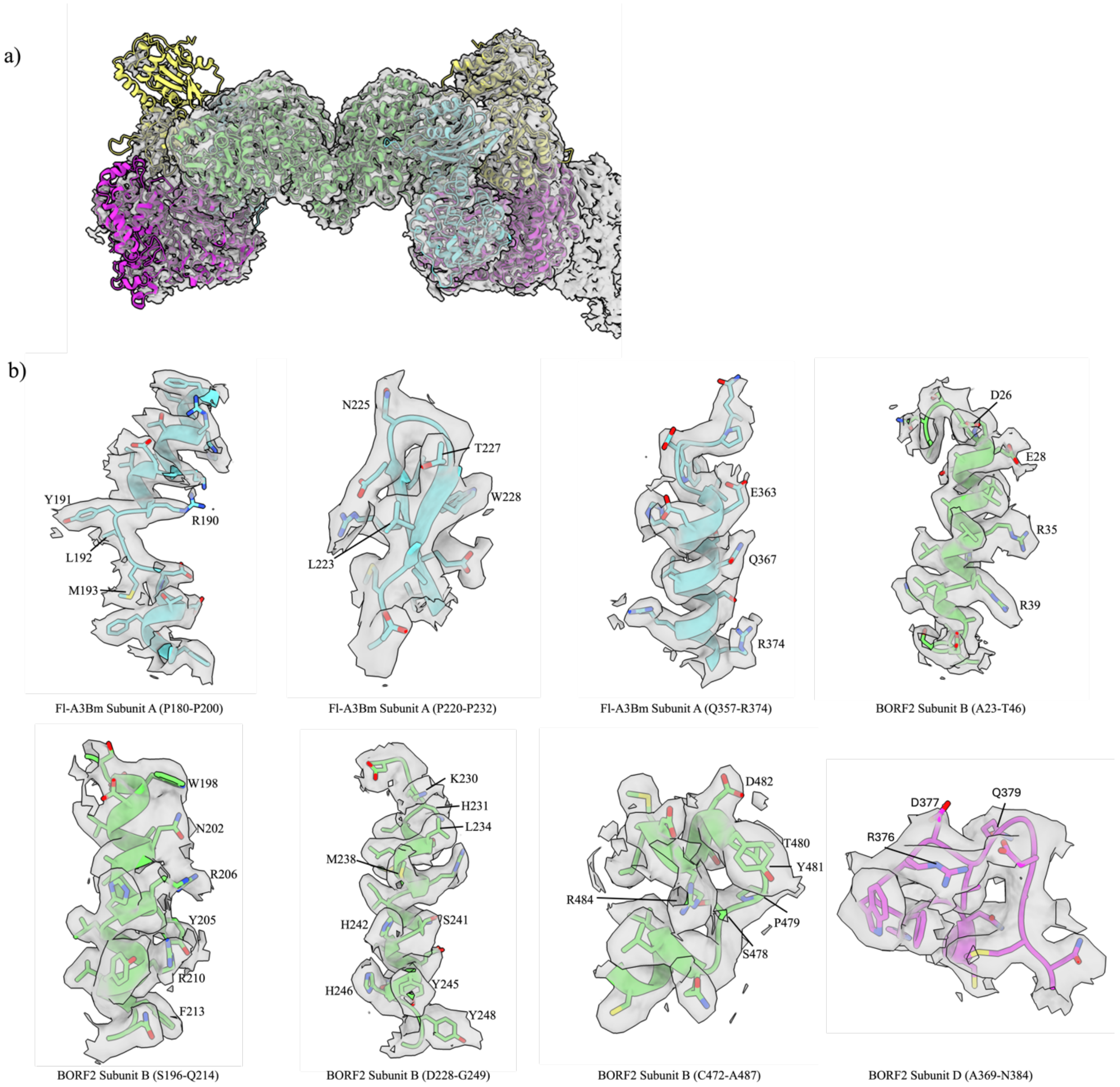


**Supplemental Figure 4. Regions of the cryo-EM density maps superposed with the atomic model of fl-A3Bm/BORF2 hetero-octamer. (a)** Global cryo-EM density of the fl-A3Bm/BORF2 hetero-octamer superimposed with its atomic model. **(b)** Segmented cryo-EM densities of representative local regions of fl-A3Bm/BORF2 are drawn as semi-transparent surfaces superposed with the corresponding atomic models of amino acid residues (ribbons and sticks), showing well-featured main-chain and side-chain density.

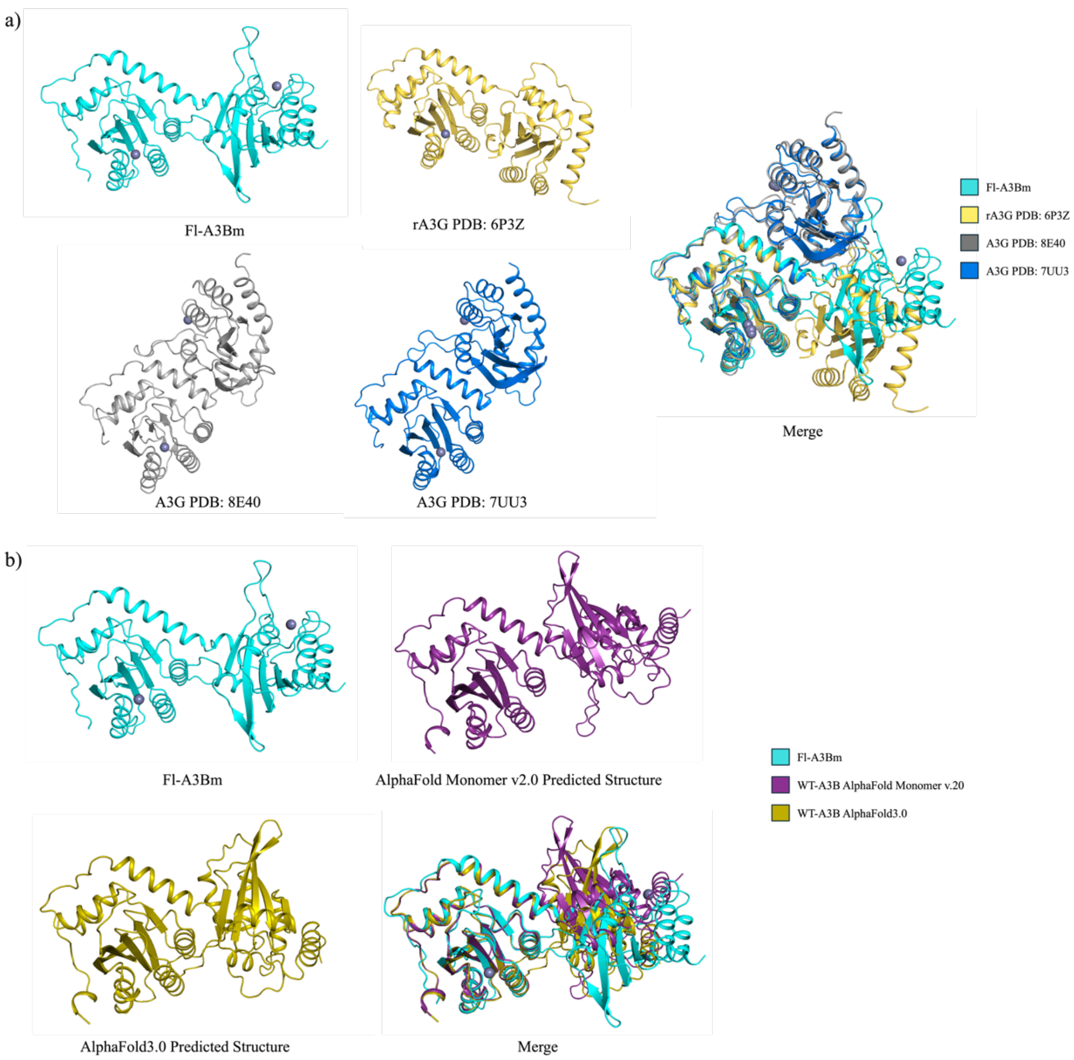


**Supplementary Figure 5. Comparison of fl-A3Bm monomer with previously published fl-double-domain A3G structures and the predicted AlphaFold A3B structures. (a)** Overlap of fl-A3Bm with previously published fl-A3G structures (PDBs 6P3Z, 8E40, 7UU3), revealing the different angles between the NTD and CTD. **(b)** Overlap of fl-A3Bm with the AlphaFold predicted A3B structures, showing different 3D domain arrangement between the NTD and CTD.

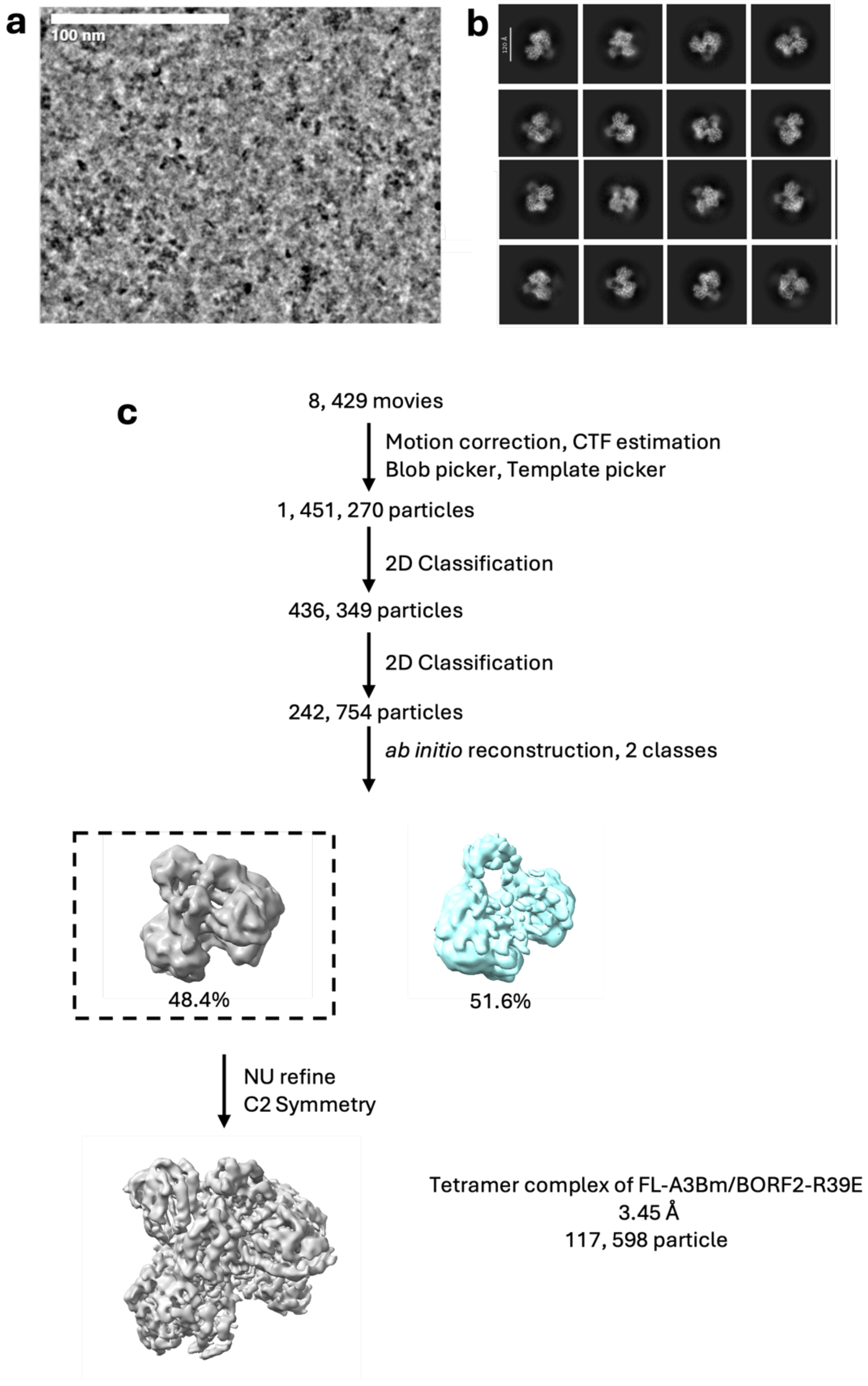


**Supplementary Figure 6. Workflow and cryo-EM 3D reconstruction of the tetrameric complex of fl-A3Bm/BORF2-R39E. (a)** Representative cryo-EM raw image. **(b)** Representative 2D class averages. **(c)** Cryo-EM image processing workflow.

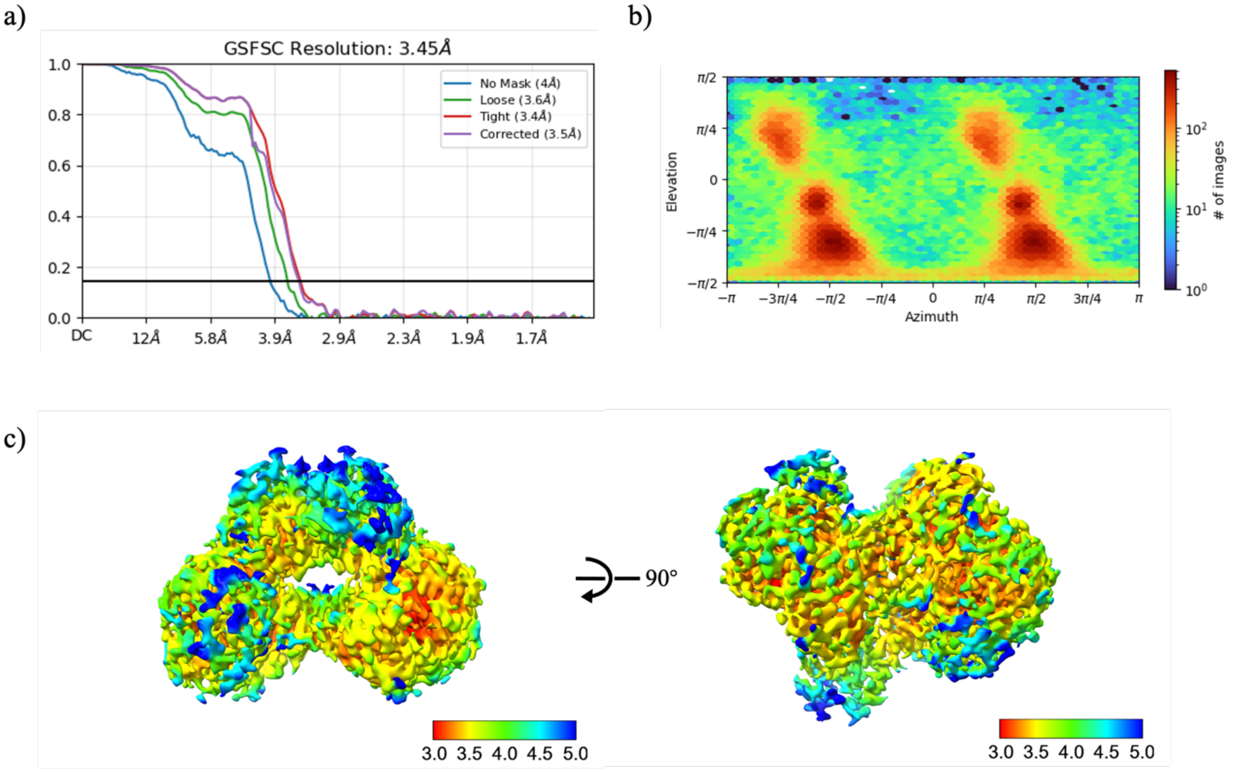


**Supplementary Figure 7. Global resolution, particle angular distribution, and local resolution estimation of the tetrameric fl-A3Bm-BORF2-R39E complex. (a)** Global resolution estimation. **(b)** Angular distribution plot of the particles. **(c)** Local resolution evaluation of the fl-A3Bm-BORF2-R39E complex.

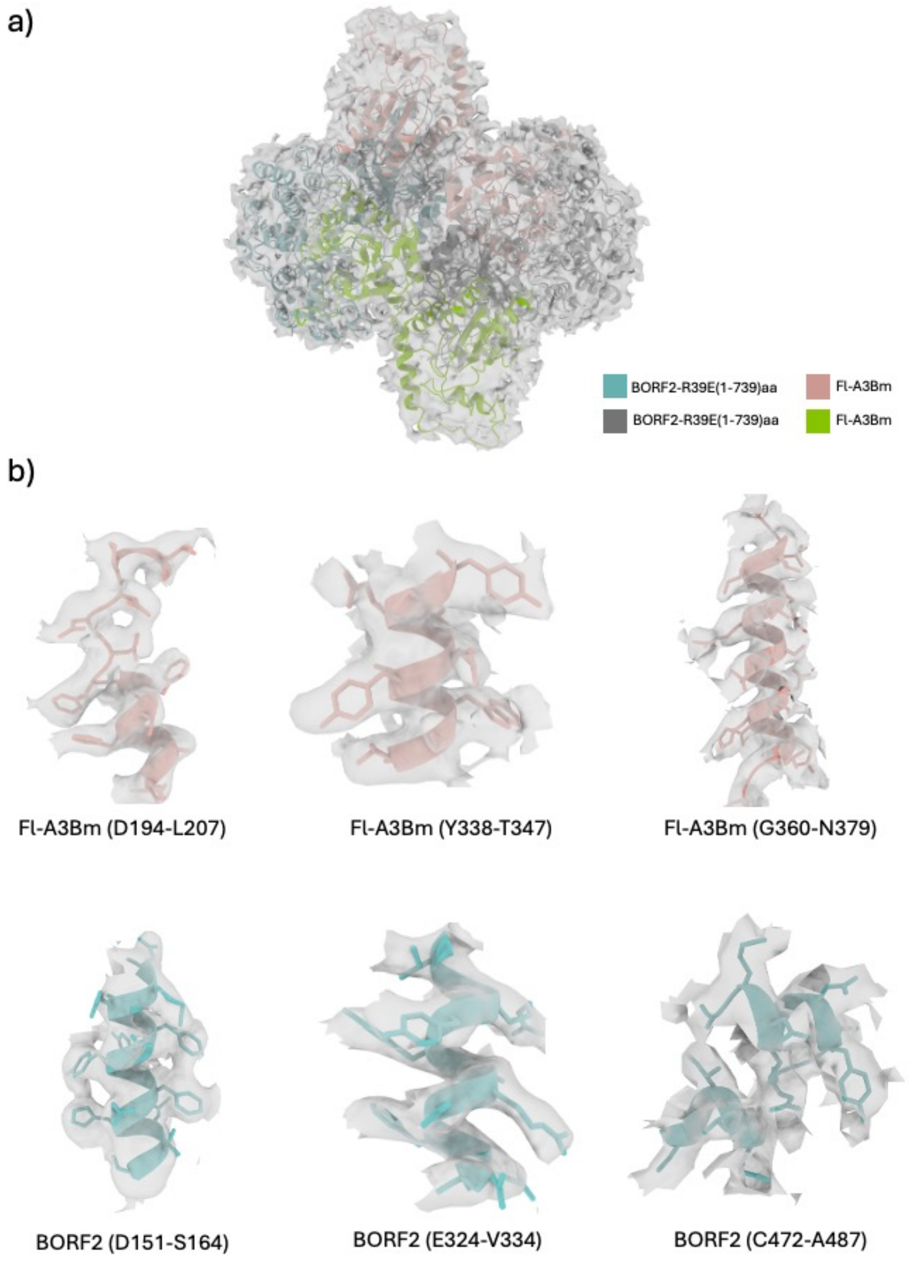


**Supplementary Figure 8. Regions of the cryo-EM density maps superimposed with the atomic model of fl-A3Bm-BORF2-R39E tetramer. (a)** Global cryo-EM density of the fl-A3Bm/BORF2-R39E tetramer superimposed with its atomic model. **(b)** Sections of cryo-EM densities of representative local regions of fl-A3Bm/BORF2-R39E complex structure are drawn as semi-transparent surfaces superimposed with the corresponding atomic models of amino acid residues (ribbons and sticks), showing well-featured main-chain and side-chain density.
